# Long-lasting impact of chito-oligosaccharide application on strigolactone biosynthesis and fungal accommodation promotes arbuscular mycorrhiza in *Medicago truncatula*

**DOI:** 10.1101/2022.08.09.503278

**Authors:** Veronica Volpe, Matteo Chialva, Teresa Mazzarella, Andrea Crosino, Serena Capitanio, Lorenzo Costamagna, Wouter Kohlen, Andrea Genre

## Abstract

- The establishment of arbuscular mycorrhiza (AM) between plants and Glomeromycotina fungi is preceded by the exchange of chemical signals: fungal released Myc-factors, including chitoligosaccharides (CO) and lipo-chitooligosaccharides (LCO), activate plant symbiotic responses, while root exuded strigolactones stimulate hyphal branching and boost CO release.
- Furthermore, fungal signaling reinforcement through CO application was shown to promote AM development in *Medicago truncatula*, but the cellular and molecular bases of this effect remained unclear.
- Here we focused on long-term *M. truncatula* responses to CO treatment, demonstrating its impact on the transcriptome of both mycorrhizal and non-mycorrhizal roots over several weeks and providing a novel insight into the mechanistic bases of the CO-dependent promotion of AM colonization.
- CO treatment caused the long-lasting regulation of strigolactone biosynthesis and fungal accommodation related genes. This was mirrored by an increase in root didehydro-orobanchol content, and the promotion of accommodation responses to AM fungi in root epidermal cells. Lastly, an advanced down-regulation of AM symbiosis marker genes was observed at the latest time point in CO-treated plants, in line with an increased number of senescent arbuscules.
- Overall, CO treatment triggered molecular, metabolic and cellular responses underpinning a protracted acceleration of AM development.

## Introduction

Mineral nutrition of most plants is supported by the mutualistic root symbiosis with Glomeromycotina, an ancient group of soil fungi (Spatafora et al., 2016) that grant their host plants preferential access to soil inorganic nutrients, in change for plant-photosynthesized sugars and lipids (Smith and Read, 2008; Wewer et al., 2014; Keymer et al., 2017).

Plant-fungus recognition is essential for AM establishment and is based on an exchange of chemical signals (Bonfante and Requena, 2011; Zipfel and Oldroyd, 2017). On the one hand, root-exuded strigolactones (SL), a class of terpenoid lactones, signal host proximity (Akiyama et al., 2005) activating hyphal metabolism and branching, and eventually promoting physical encounter with the root surface (Besserer et al., 2006; Waters et al., 2017). On the other hand, AM fungi release diffusible molecules (Myc-factors) that activate a so-called Common Symbiotic Signalling Pathway, or CSSP (Zipfel and Oldroyd, 2017; Choi et al., 2018). Downstream responses include local and systemic changes in gene expression and metabolism, overall preparing the host plant to symbiosis establishment (MacLean et al., 2017; Choi et al., 2018; Pimprikar and Gutjarh, 2018). Myc-factors include two classes of molecules: lipo-chito-oligosaccharides, or LCO (Maillet et al., 2011), structurally similar to rhizobial Nod-factors and composed of a short chitin chain with a few lateral substitutions; and short-chain chito-oligosaccharides, or CO (Genre et al., 2013), where only the chitin backbone is present. The activity of CO as AM fungal signals has been demonstrated in all tested host plants, including monocots and dicots (Nasir et al., 2021) and their release is boosted upon strigolactone perception (Genre et al., 2013). Furthermore, CO can easily be produced through chitin hydrolysis (Crosino et al., 2021) making them particularly interesting for large scale agricultural applications (Volpe et al., 2020). In this context, we recently demonstrated that plant treatment with exogenous CO prior to fungal inoculation promotes AM colonization with a marked increase in arbuscule development, biomass accumulation and total photosynthetic surface, compared to untreated mycorrhizal plants (Volpe et al., 2020).

Most of the published research has focused on short-term plant responses to CO (or LCO) treatment, demonstrating the activation of symbiotic signaling and gene regulation in the range of a few hours (Czaja et al., 2012; Camps et al., 2015; Giovannetti et al., 2015; Feng et al., 2019). Here we chose to focus on a longer time scale, in an attempt to track the molecular bases of the CO-dependent promotion of AM development over several weeks (Volpe et al., 2020). To this aim, we used RNA-seq to investigate genome-wide changes in root gene expression over four weeks after the initial short-chain CO treatment, and further validated our results with functional insights. CO treatment changed the expression pattern of the whole strigolactone biosynthetic pathway and increased the didehydro-orobanchol content in root tissue. Moreover, RNA-seq data, targeted gene expression analyses and live imaging of epidermal cell reorganization indicated a CO-dependent stimulation of intracellular accommodation processes. Lastly, the down-regulation of AM symbiosis marker genes was consistent with the increased number of senescent arbuscules at the end of our experimental time frame. In conclusion, our results revealed that CO treatment impacted on molecular, cellular and metabolic mechanisms that converged towards a global acceleration in AM development.

## Materials and Methods

### Plant and fungal materials

The model legume *Medicago truncatula* cv ‘Jemalong’ (line A17) was used for this study. Seeds were collected from the pods, conserved for 2 days at room temperature to allow the oxygenation of internal tissues and scarified on sandpaper in order to break the seed coat. Seeds were then sterilized using 5% (v/v) sodium hypochlorite in sterile water and rinsed in sterile distilled water, each step for 5 min keeping the falcon tubes under constant stirring. To break dormancy and allow rapid and synchronized germination, seeds were placed on 0.6% Plant-Agar (Duchefa, Haarlem, The Netherlands) plates, maintained at 4°C in the dark for two days and then moved at 23°C until germination.

A commercial granular inoculum of the AM fungus *Funneliformis mosseae* (strain BEG 12) from MycAgroLab (www.mycagrolab.com; France) was used for all experiments. This inoculum was composed by the growth substrate of *Sorghum vulgare* plants, colonized root pieces, spores and extraradical mycelium, with a minimum of 10 active propagules per g of inoculum.

### Chitin oligomers

A mixture of short-chain chito-oligosaccharides, CO (Volpe et al. 2020) was used for all plant treatments. The CO mixture was obtained from crustacean food manufacturing waste (Zhengzhou Sigma Chemical Co., Ltd. Zhengzhou, Henan, China) and contained fully acetylated, mono-deacetylated and di-deacetylated molecules composed of 2 to 5 N-acetyl-glucosamine residues (Volpe et al. 2020). CO were applied as a 1 g/L solution in sterile distilled water supplemented with Tween 20 (0.005%) as a surfactant. Sterile water with Tween 20 (0.005%) was used in control treatments.

### Pot culture

Young *M. truncatula* seedlings were transferred into 9 cm x 9 cm x 12 cm pots (one seedling per pot) filled with sterile coarse sand (0.4-0.8 mm; Valle Po, Revello, CN, Italy). A plastic bag was placed on the pots during the first week, to protect the young plants from desiccation. Plants were grown in phytochambers under controlled conditions (23°C day/21°C night temperature, 16/8 h light/dark photoperiod) and fertilized once per week using a modified Long Ashton (LA) nutrient solution (Hewitt, 1966) containing 3.2 μM phosphate and 1mM nitrate. During the rest of the week, pots were watered when necessary with tap water. Four experimental conditions were set up: control (CTR) plants, lacking both CO treatment and AM inoculation; CO-treated control (CTR+CO); mycorrhizal (MYC), where AM inoculum was added in the absence of CO treatments; and CO-treated mycorrhizal (MYC+CO), where plants were both inoculated and exposed to the CO solution. CO treatment and fungal inoculum were applied as previously described in Volpe et al. (2020). In short, 5 ml of CO solution were sprayed over the pot substrate surface 4 and 2 days before inoculation with 22.5 ml of fungal inoculum.

Plants were harvested at 10, 14, 21, 28 days post-inoculation (dpi) and carefully washed under tap water to remove sand. AM colonization was assessed based on the presence of root-bound extraradical mycelium prior to rapid sampling, dip-freezing in liquid nitrogen and storage at −80°C until RNA extraction. We used 5 plants for each experimental condition at each time point, except at 10 dpi, when the limited extension of root systems required the sampling of a larger number of plants (12). We collected at least three biological replicates (plants) for each condition, selecting the best developed plants, with no visible sign of pathogenesis or stress and an extensive extraradical fungal development.

### RNA isolation and sequencing

Total RNA was extracted from all samples using the RNeasy^™^ Plant Mini kit (Qiagen, Hilden, Germany). Samples were mechanically homogenized using a TissueLyser system for 2 minutes at 18 Hz. An aliquot of each sample was mixed with RLT buffer (Qiagen), and further treated following the manufacturer’s protocol. RNA quantity and quality were spectrophotometrically checked using a NanoDrop ND-1000 instrument. RNA samples were further quantified and tested for integrity by capillary electrophoresis using Agilent 2100 Bioanalyzer instrument with the Agilent RNA 6000 Nano Kit following manufacturer’s instructions. As requested by the sequencing company (IGATech, Udine, Italy) samples dedicated to RNA-seq were not treated with DNase.

RNA-seq analysis on root RNA samples was performed at IGA Technology services. TruSeq stranded mRNA kit (Illumina, San Diego, CA) was used for library preparation and sequencing performed using Illumina NextSeq 500 platform (Illumina, San Diego, CA) in single-end mode at 75 bp read-length and with a sequencing depth ranging from 20 to 30 M of reads per sample (Table S1).

### Bioinformatics

Adapter sequences were masked with Cutadapt v1.11 (Martin, 2011) from raw reads using the following parameters: --anywhere (on both adapter sequences) --overlap 5 -times 2 –minimum-length 35-mask-adapter. Raw reads were then trimmed removing lower quality bases and adapters using ERNE software (Del Fabbro et al., 2013). The expression level of each gene in each library was calculated by mapping filtered reads on Mt4.1 reference genome (Young et al., 2011; Tang, et al., 2014) using STAR splice-aware aligner (Dobin et al., 2013) with default parameters. Reads overlapping with annotated exons were counted using HTSeq (Anders et al., 2015) and differential expression analysis performed for each comparison (CTR+CO *vs* CTR, MYC vs CTR, MYC+CO *vs* CTR and MYC+CO *vs* MYC) at each time points (10, 14, 21,28 dpi) using DESeq2 R package (Love et al., 2014) at a False discovery rate (FDR or adjusted p-value) (Benjamini et al., 1995) threshold of 0.05 (SI_DATA1-4). The RNA-seq dataset obtained in this study was included in the Medicago Expression Atlas (MtExpress V3, Carrere et al., 2021).

Differentially expressed genes were plotted into KEGG (Kyoto Encyclopedia of Genes and Genomes) pathway maps using the ‘pathview’ R package (Luo et al., 2013) and the *M. truncatula* annotation available at the KEGG website. Metabolic pathways visualization was achieved using MapMan software (Usadel et al., 2005). MapMan functional annotation for *M. truncatula* (Table S2) was obtained from the proteome using Mercator4 annotation tool (Schwacke et al., 2019).

Gene-ontology (GO) enrichment analyses were carried out on each of the previously generated contrasts using GOseq R package (Young et al., 2010) at a FDR cutoff of 0.1. The −log_10_ of the corrected p-value was plotted.

Variance Partitioning Analysis (VPA) was performed using the ‘varpart’ function in R package ‘vegan’ (Oksanen et al., 2019) using gene counts normalized with DESeq2 through the Variance Stabilizing Transformation (vst). Fractions of variance explained by single factors were tested for significance on the RDA model using permutational ANOVA (999 permutation, p < 0.05). Principal component analysis (PCA) was calculated with the ‘prcomp’ function in ‘base’ R package, using same normalization used for VPA. Statistical and graphical elaborations were performed in R programming environment (R Core Team 2020) using ‘ggplot2’ package (Wickham et al., 2022).

### Gene expression analysis by real-time qPCR

RNA-seq data was validated by quantitative real-time PCR (qRT-PCR) analysis of the expression profile of 10 differentially-expressed genes (DEGs) across all treatments and time points (Fig. S1). Specific primers (listed in Table S3) were designed using PerlPrimer software (http://perlprimer.sourceforge.net) on *M. truncatula* CDS sequences from NCBI (http://www.ncbi.nlm.nih.gov/) and *M. truncatula* genome database (http://www.medicagogenome.org). All primers were first tested by PCR on genomic *M. truncatula* and *F. mosseae* DNA, to confirm their plant-specificity.

To remove any trace of genomic DNA before cDNA synthesis, RNA samples were treated with TURBO^™^ DNase (QIAGEN, Hilden, Germany) according to the manufacturer’s instructions, followed by second NanoDrop quantification. The RNA samples were routinely checked for DNA contamination by PCR analysis, using primers *MtTEF-F:* 5’-AAGCTAGGAGGTATTGAAAG- 3’ and *MtTEF-R:* 5’- ACTGTGCAGTAGTACTTGGTG-3’ for *MtTEF* (Elongation Factor, NCBI Ref. Seq.: XM_013595882).

For single-strand cDNA synthesis, approximately 700 ng of total RNA was denatured at 65°C for 5 min and then reverse-transcribed using the Super-Script II kit (Invitrogen, Carlsbad, CA, USA) at 25°C for 10 min, 42°C for 50 min and 70°C for 15 min. The final volume was 20 μl volume and contained 10 μM of random primers, 0.5 mM deoxynucloside triphosphates, 4 μl 5x buffer, 2 μl 0.1M dithiothreitol (DTT) and 1 μl Superscript II enzyme Control PCR was set with TEF primers to test the presence of cDNA.

To confirm the results of our transcriptomic analysis of SL biosynthesis and transport related genes, root systems from CTR and CTR+CO plants were sampled at 21 dpi in an independent experiment. 100 mg of root powder was used for RNA extraction and analyzed by qRT-PCR.

Lastly, in order to monitor local, short-time occurrence of early gene regulation in response to CO perception, 1 g/L, 1 mg/L or 1 μg/L CO solutions were applied to WT and *dmi3-1 Agrobacterium rhizogenes*-transformed root organ cultures (ROCs) expressing the nuclear targeted NUPYC2.1 protein (Sieberer et al., 2009), which was used as a visual reporter of cell viability. ROC lines were propagated on M medium (Bècard & Fortin, 1988) at 25°C in the dark, in vertically oriented petri dishes to favor the regular fishbone-shaped root system (Chabaud et al., 2002) and grown for 3 weeks. For local CO treatment, small discs of filter paper (Ø 4mm) soaked in 20 μl of CO solution were applied on several lateral roots, 10-15 mm from the root tip, in agreement with Chabaud et al. (2011). Six hours later, 1 cm-long root segments underlying each disc were excised, immediately frozen and stored at −80°C until RNA extraction and analysis for the expression of AM fungal perception and accommodation marker genes.

qRT-PCR analyses were done according to Rotor-Gene SYBR Green PCR Kit instructions (QIAGEN, Hilden, Germany) and were run in a final volume of 15 μl for each tube containing 7.5 μl of Rotor-Gene SYBR Green PCR Master Mix, 5.5 μl of 3 μM specific primers mix (Table S4), and 10 ng of cDNA. A Rotor Gene machine (Qiagen) was used with the following program: 3 min pre-incubation at 95°C, followed by 40 cycles of 15s at 95°C, and 30s at 60°C. Each amplification was followed by melting curve analysis (60–94°C) with a heating rate of 0.5°C every 15s. All reactions were performed with two technical replicates and only Ct values with a standard deviation that did not exceed 0.5 were considered. Relative RNA levels were calibrated using the elongation factor (TEF) mRNA as endogenous reference and normalized to the control line. Results were validated statistically using the unpaired Student’s t-test to compare two means: differences were considered significant at P<0.05.

### Strigolactone analysis in root systems and exudates

CTR and CTR+CO plants were twice respectively treated with water or CO solution, and then grown for 21 days. 21 day-old *M. truncatula* plants were carefully sampled and washed free of sand. They were put inside a tube (2 plants for tube) with 40 mL of LA solution covering the whole root system and closed with Parafilm; the tubes were placed into a closed Magenta box and the root system was covered with a black shield. Plants (4 biological replicates for each treatment) were placed in a growth chamber for 36 h for exudate collection. After 36 hours, the root exudate was recovered and stored at −20°C for the next step.

Whole plant fresh weight, as well as shoot and root biomass separately, were measured for each biological replicate, before each root system was ground to a fine powder in a mortar using liquid nitrogen and stored at −80°C.

For the analysis of SL content, 500 mg of each ground sample were transferred to 2 mL eppendorf tubes and SL were extracted with 2 mL of ethyl acetate with 10^-8^ M GR24 as internal standard (end concentration 10^-7^ M). Tubes were vortexed, sonicated for 20 sec (sonication bath, Branson Ultrasonic) and centrifuged for 10 min at 10,000 rpm at RT. Subsequently, the organic phases were transferred to 4 mL glass vials and the solvent was evaporated in a speed vacuum system (SPD121P, ThermoSavant, Hastings, UK). Root exudates were purified and concentrated as previously described for tomato (Kohlen et al., 2012); SL from root tissues and exudates were quantified according to Liu et al. (2011).

### Live imaging of prepenetration responses

To further investigate whether CO treatment altered cell responses to fungal contact, we modified the targeted AM inoculation technique used by Genre et al. (2005; 2008) for visualizing the pre-penetration apparatus (PPA) in WT roots. In short, *Agrobacterium rhizogenes*-transformed root organ cultures (ROCs) expressing the endoplasmic reticulum-targeted GFP-HDEL (Haseloff et al., 1997) were grown on M medium (Boisson-Dernier et al., 2001) in vertically-oriented Petri dishes and inoculated with pre-germinated spores of *Gigaspora margarita*. 1 ml of filtered CO solution (1 mg/L) or sterilized water (as control) was then added over the root culture before covering it with a gas-permeable plastic film (bioFOLIE 25; Sartorius, Goettingen, Germany). Fungus-root interaction was monitored daily using a stereomicroscope and contact sites were imaged at 7, 10 and 14 dpi using a Leica TCS SP2 confocal microscope fitted with a long distance 40X water-immersion objective (HCX Apo 0.80). GFP fluorescence was excited with the argon laser band at 488 nm and recorded with an emission window set at 500-525 nm.

### Morphological and functional analysis of AM colonization

Total and root fresh biomass were measured in mycorrhizal WT plants in the presence or absence of CO treatment. Biomass data were compared and the results were validated statistically using the unpaired student’s t-test (differences were considered significant at P<0,05). Furthermore, inoculated WT and *dmi3-1* mutant plants, treated or not with CO, were sampled at 28 dpi to quantify fungal colonization according to Trouvelot et al. (1986). At least four plants were used for the root mycorrhization intensity assessment and 100 1 cm-long root pieces were analyzed per plant. The same method was adapted for the *dmi3-1* mutant (where intraradical colonization is blocked) to determine frequency (F) and intensity (M) parameters with reference to hyphopodium presence on the root surface instead of intraradical fungal structures.

For detailed microscope analysis of arbuscule morphology, root segments of WT plants were excised and individually embedded in agarose (5%). 100 μm vibratome sections were then moved to microscope slides and treated for 5 min in phosphate buffer containing 0.5% commercial bleach, rinsed three times, and then incubated overnight in 10 μg/mL wheat germ agglutinin-fluorescein isothiocyanate (WGA-FITC; Sigma-Aldrich, Milan, Italy) to label the fungal wall. A Leica TCS SP2 confocal microscope (Leica Microsystems GmbH, Wetzlar, Germany) equipped with a 40X water immersion objective was used for imaging with fluorescence excitation at 488 nm and acquisition at 500-550 nm.

## Results

### CO treatment impacted on the root transcriptome for several weeks

In a first global analysis of the RNA-seq dataset, principal component analysis (PCA; Fig. 1a) and variance partitioning analysis (VPA; Fig. 1b) revealed a high level of variation in the transcriptome between conditions and time points. As shown in Figure 1a, CTR, CTR+CO, MYC and MYC+CO transcriptomes clustered in different areas of the PCA plot, indicating that both AM inoculation and CO treatment had a strong influence on the root gene expression profiles. In order to grant a homogeneous code for sample timing, we refer to dpi (days post inoculation) for both inoculated and non-inoculated (but same age) plants. At 10 dpi, CTR+CO, MYC and MYC+CO samples were clearly separated from CTR along the PC1 axis (explaining 33.38% of variance), highlighting the distance in the transcriptional profile of CTR samples compared to the remaining treatments. A major distinction along the PC1 axis became evident since 14 dpi between inoculated (MYC and MYC+CO) and non-inoculated samples (CTR and CTR+CO), as expected in relation with AM establishment. A specific effect of the CO treatment was evident at 10 dpi (with a clear separation between CTR and CTR+CO, and a milder but evident separation between MYC and MYC+CO), and partially rebounded at 28 dpi between MYC and MYC+CO samples.

**Fig. 1.**
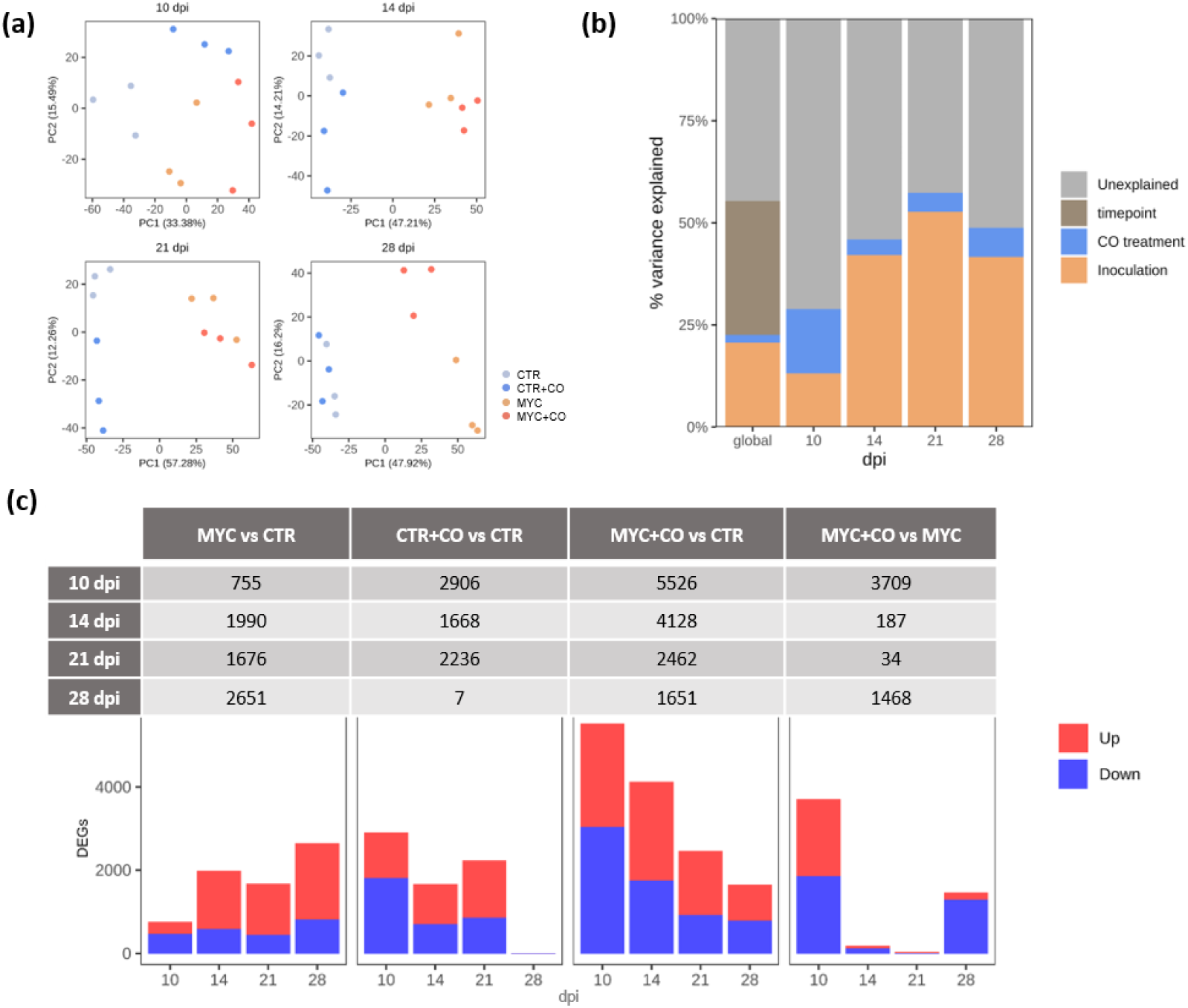
Global overview of transcriptomic data. **(a)** Principal component analysis (PCA) plots highlighted a clear separation of the expression profiles between AM inoculated (dark and light orange) and non-inoculated (dark and light blue) samples, as well as between treated (dark orange e dark blue) and non-treated (light orange and light blue) ones. Three biological replicates (individual dots) are plotted per condition and time point. **(b)** Variance Partitioning Analysis (VPA) considered three factors: AM fungus presence (inoculation), CO treatment and time (10, 14, 21,28 dpi). The strongest influence on gene regulation resulted to depend on time, while CO treatment had a major effect at the earliest time point (10 dpi). All the fractions explained by the factors were significant (ANOVA on RDA model, p<0.05), except for the fraction explained by CO treatment at 14 dpi. **(c)** Differentially expressed genes (DEGs) in the four comparisons and at different time points. Number and expression trend of DEGs. Up-regulation is shown in red, down-regulation in blue (FDR <0.05).

VPA analysis (Fig. 1b) highlighted time as the most influential variable in our time course study, in line with the expected major changes in gene expression throughout AM development (Handa et al., 2015). VPA was subsequently used to address individual time points: this correlated CO treatment to the highest amount of variance in gene regulation at 10 dpi, an early time point in root colonization in our experimental conditions, while AM inoculation became the most influential variable in the later time points. Also in this case the CO effect extended across all four time points, with a partial reinforcement at 28 dpi.

In conclusion, both PCA and VPA indicated an impact of CO treatment across our whole time course analysis, with the strongest effect at the earliest time point in both inoculated and non-inoculated plants.

The number and proportion of differentially expressed genes (DEGs) among the four conditions at each of the four time points is summarized in Figure 1c. The MYC *vs* CTR comparison highlighted a progressive increase in gene regulation, in line with the ongoing colonization of the root system by the AM fungus. In the CTR+CO *vs* CTR comparison, the CO effect was strongest at 10 dpi (with 2906 DEGs), intermediate at 14 and 21 dpi, and minimal at 28 dpi (with only 7 DEGs). This pattern can be explained with the progressive degradation/leaching of CO from the pot over the weeks that follow the initial treatments (4 and 2 days before inoculation). The strongest differences in gene regulation were observed when comparing MYC+CO *vs* CTR samples (which represent the most dissimilar conditions in our experiment). Here, the strongest impact of the CO treatment was recorded at 10 dpi (5526 DEGs), with a progressive decrease thereafter. This is suggestive of a synergistic effect of exogenous CO and fungal-secreted Myc-factors during early root colonization, and is in line with the progressive decrease of CO effect over time (as highlighted in CTR+CO *vs* CTR). Lastly, the MYC+CO *vs* MYC comparison aimed at dissecting the effect of exogenous CO on the AM interaction. Also in this case, a major impact was evident at 10 dpi (3709 DEGs), in line with the reinforcement of fungal signaling by the applied CO. Nevertheless, differential gene regulation peaked again at 28 days (1468 DEGs) with a large majority of down-regulated sequences, whereas intermediate time points displayed a very similar gene regulation scenario in the two conditions.

In summary, differential gene expression analysis convincingly supported the evidence emerging from multivariate analysis, indicating an impact of CO on plant gene regulation throughout our experimental time frame, with a major effect during early root colonization.

### Functional enrichment analysis suggests that CO treatment mimics fungal presence

To gain insights into the molecular processes impacted by CO treatment, we performed a Gene Ontology (GO) enrichment analysis of DEGs (SI_DATA5-8; http://geneontology.org/docs/ontology-documentation/). We will focus here on the two most strongly regulated time points: 10 and 28 dpi, and the most over-represented GO terms for each comparison, as presented in Figure S2-S5. Interestingly, in the MYC *vs* CTR comparison (Fig. S2) *Defense responses* were over-represented both at 10 and 28 dpi, in agreement with the well documented regulation of defense-related genes during AM interaction (Salzer et al., 2000; Jung et al., 2012; Cameron et al., 2013). *Protein serine/threonine kinase activity* emerged at 10 and 28 dpi and *DNA-binding transcription factor activity* at 28 dpi, reflecting the extensive impact of symbiosis on regulatory pathways (Liu et al., 2003; Küster et al., 2004; Sanchez et al., 2004; Hohnjec et al., 2005; Hogekamp and Küster, 2013). The whole ubiquitination system (*ubiquitin ligase complex, protein ubiquitination, ubiquitin protein transferase activity*) was regulated at 28 dpi, possibly indicating the onset of senescence-related mechanisms. Lastly, *fatty acid biosynthesis* appeared at 28 dpi, likely related to both the extensive synthesis of perifungal membranes and the intense production of lipids feeding the fungus (Wewer et al., 2014; Luginbuehl et al., 2017; MacLean et al. 2017).

Since no enriched GO category emerged at 28 dpi for the CTR+CO *vs* CTR comparison (Fig. S3), we only comment here on the 10 dpi time point. In this case, a remarkable analogy emerged with the same time point of the MYC *vs* CTR comparison (Fig. S6): in fact, the global pattern of gene regulation was very similar for *kinase activity, signal transduction, defense response* and *acyl transferase activity*. This supports the role of CO as mimics of fungal presence and elicitors of symbiotic signaling and gene regulation in the host root.

In the MYC+CO *vs* CTR comparison at 10 dpi (Fig. S4), a massive activation was observed in protein biosynthesis (*translation, ribosome, ribosome constituents*), cytoskeleton-associated processes (*microtubule associated complex, microtubule-based movement and microtubule motor activity*) and chromatin reorganization (*nucleosome, nucleosome assembly*). This is suggestive of intense cell reorganization and is supported by our knowledge of the fungal accommodation process that takes place inside each colonized cell (Gutjahr and Parniske, 2013; Carotenuto et al., 2019). At 28 dpi (Fig. S4), GO terms related to transport (*transporter activity*, *transport*, *heme binding*, *sulfate transport*, *transmembrane transport*, *secondary active sulfate transmembrane transport activity*, *membrane*) were highly represented, likely related to the extensive AM colonization promoted by CO treatment (Benedito et al., 2010; Gaude et al., 2012; Handa et al., 2015).

Finally, the MYC+CO *vs* MYC comparison (Fig. S5) suggested that exogenous CO enhanced *cell cycle regulation*, *chromatin rearrangement* and *cytoskeleton-associated processes* at 10 dpi, hinting at an intensification of the fungal accommodation responses in CO-treated plants. By contrast, a major impact on regulatory and proteolytic processes appeared at 28 dpi in CO-treated plants, suggesting the appearance of senescence-related processes.

In conclusion, our analysis of GO enrichment indicated a global consistency between the regulatory responses induced by AM fungi and CO treatment, with a reinforcement and an advancement of symbiotic processes in plants that were exposed to both stimuli.

### CO treatment stimulated strigolactone signaling and fungal accommodation

Among the various gene pathways related to AM establishment that were highlighted by GO enrichment analysis (Table 1), we chose to validate the observed regulation of signaling- and cell reorganization-related genes with functional analyses.

**Table 1.**
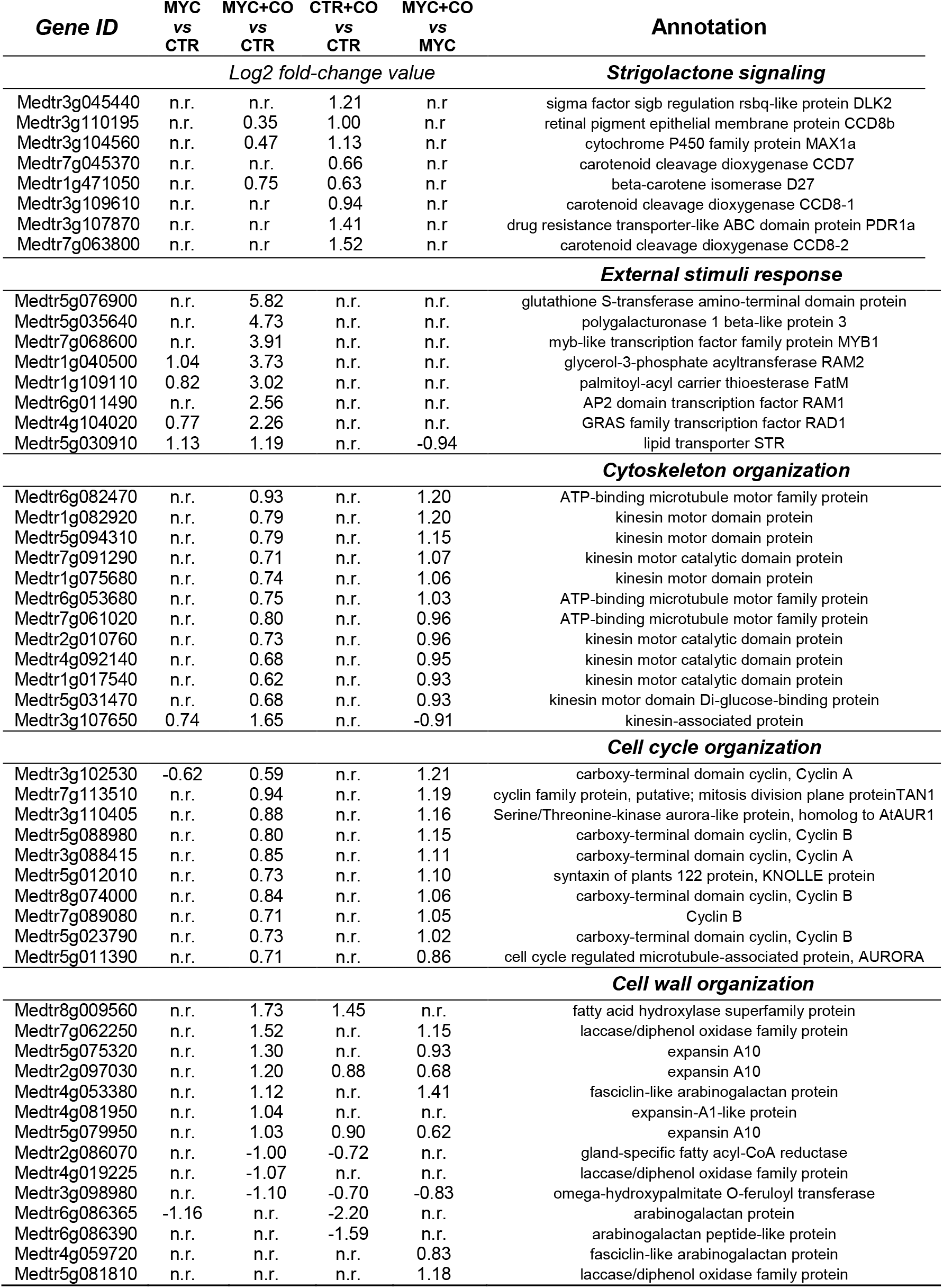
Main AM symbiosis-related gene categories impacted at 10 dpi by CO treatment (n.r.= not significantly regulated).

| *Gene ID* | MYC<br>vs<br>CTR | MYC+CO<br>vs<br>CTR | CTR+CO<br>vs<br>CTR | MYC+CO<br>vs<br>MYC | Annotation |
| --- | --- | --- | --- | --- | --- |
| *Log2 fold-change value* |  |  |  |  | ***Strigolactone signaling*** |
| Medtr3g045440 | n.r. | n.r. | 1.21 | n.r. | sigma factor sigb regulation rsbq-like protein DLK2 |
| Medtr3g110195 | n.r. | 0.35 | 1.00 | n.r. | retinal pigment epithelial membrane protein CCD8b |
| Medtr3g104560 | n.r. | 0.47 | 1.13 | n.r. | cytochrome P450 family protein MAX1a |
| Medtr7g045370 | n.r. | n.r. | 0.66 | n.r. | carotenoid cleavage dioxygenase CCD7 |
| Medtr1g471050 | n.r. | 0.75 | 0.63 | n.r. | beta-carotene isomerase D27 |
| Medtr3g109610 | n.r. | n.r. | 0.94 | n.r. | carotenoid cleavage dioxygenase CCD8-1 |
| Medtr3g107870 | n.r. | n.r. | 1.41 | n.r. | drug resistance transporter-like ABC domain protein PDR1a |
| Medtr7g063800 | n.r. | n.r. | 1.52 | n.r. | carotenoid cleavage dioxygenase CCD8-2 |
|  |  |  |  |  | ***External stimuli response*** |
| Medtr5g076900 | n.r. | 5.82 | n.r. | n.r. | glutathione S-transferase amino-terminal domain protein |
| Medtr5g035640 | n.r. | 4.73 | n.r. | n.r. | polygalacturonase 1 beta-like protein 3 |
| Medtr7g068600 | n.r. | 3.91 | n.r. | n.r. | myb-like transcription factor family protein MYB1 |
| Medtr1g040500 | 1.04 | 3.73 | n.r. | n.r. | glycerol-3-phosphate acyltransferase RAM2 |
| Medtr1g109110 | 0.82 | 3.02 | n.r. | n.r. | palmitoyl-acyl carrier thioesterase FatM |
| Medtr6g011490 | n.r. | 2.56 | n.r. | n.r. | AP2 domain transcription factor RAM1 |
| Medtr4g104020 | 0.77 | 2.26 | n.r. | n.r. | GRAS family transcription factor RAD1 |
| Medtr5g030910 | 1.13 | 1.19 | n.r. | -0.94 | lipid transporter STR |
|  |  |  |  |  | ***Cytoskeleton organization*** |
| Medtr6g082470 | n.r. | 0.93 | n.r. | 1.20 | ATP-binding microtubule motor family protein |
| Medtr1g082920 | n.r. | 0.79 | n.r. | 1.20 | kinesin motor domain protein |
| Medtr5g094310 | n.r. | 0.79 | n.r. | 1.15 | kinesin motor domain protein |
| Medtr7g091290 | n.r. | 0.71 | n.r. | 1.07 | kinesin motor catalytic domain protein |
| Medtr1g075680 | n.r. | 0.74 | n.r. | 1.06 | kinesin motor domain protein |
| Medtr6g053680 | n.r. | 0.75 | n.r. | 1.03 | ATP-binding microtubule motor family protein |
| Medtr7g061020 | n.r. | 0.80 | n.r. | 0.96 | ATP-binding microtubule motor family protein |
| Medtr2g010760 | n.r. | 0.73 | n.r. | 0.96 | kinesin motor catalytic domain protein |
| Medtr4g092140 | n.r. | 0.68 | n.r. | 0.95 | kinesin motor catalytic domain protein |
| Medtr1g017540 | n.r. | 0.62 | n.r. | 0.93 | kinesin motor catalytic domain protein |
| Medtr5g031470 | n.r. | 0.68 | n.r. | 0.93 | kinesin motor domain Di-glucose-binding protein |
| Medtr3g107650 | 0.74 | 1.65 | n.r. | -0.91 | kinesin-associated protein |
|  |  |  |  |  | ***Cell cycle organization*** |
| Medtr3g102530 | -0.62 | 0.59 | n.r. | 1.21 | carboxy-terminal domain cyclin, Cyclin A |
| Medtr7g113510 | n.r. | 0.94 | n.r. | 1.19 | cyclin family protein, putative; mitosis division plane proteinTAN1 |
| Medtr3g110405 | n.r. | 0.88 | n.r. | 1.16 | Serine/Threonine-kinase aurora-like protein, homolog to AtAUR1 |
| Medtr5g088980 | n.r. | 0.80 | n.r. | 1.15 | carboxy-terminal domain cyclin, Cyclin B |
| Medtr3g088415 | n.r. | 0.85 | n.r. | 1.11 | carboxy-terminal domain cyclin, Cyclin A |
| Medtr5g012010 | n.r. | 0.73 | n.r. | 1.10 | syntaxin of plants 122 protein, KNOLLE protein |
| Medtr8g074000 | n.r. | 0.84 | n.r. | 1.06 | carboxy-terminal domain cyclin, Cyclin B |
| Medtr7g089080 | n.r. | 0.71 | n.r. | 1.05 | Cyclin B |
| Medtr5g023790 | n.r. | 0.73 | n.r. | 1.02 | carboxy-terminal domain cyclin, Cyclin B |
| Medtr5g011390 | n.r. | 0.71 | n.r. | 0.86 | cell cycle regulated microtubule-associated protein, AURORA |
|  |  |  |  |  | ***Cell wall organization*** |
| Medtr8g009560 | n.r. | 1.73 | 1.45 | n.r. | fatty acid hydroxylase superfamily protein |
| Medtr7g062250 | n.r. | 1.52 | n.r. | 1.15 | laccase/diphenol oxidase family protein |
| Medtr5g075320 | n.r. | 1.30 | n.r. | 0.93 | expansin A10 |
| Medtr2g097030 | n.r. | 1.20 | 0.88 | 0.68 | expansin A10 |
| Medtr4g053380 | n.r. | 1.12 | n.r. | 1.41 | fasciclin-like arabinogalactan protein |
| Medtr4g081950 | n.r. | 1.04 | n.r. | n.r. | expansin-A1-like protein |
| Medtr5g079950 | n.r. | 1.03 | 0.90 | 0.62 | expansin A10 |
| Medtr2g086070 | n.r. | -1.00 | -0.72 | n.r. | gland-specific fatty acyl-CoA reductase |
| Medtr4g019225 | n.r. | -1.07 | n.r. | n.r. | laccase/diphenol oxidase family protein |
| Medtr3g098980 | n.r. | -1.10 | -0.70 | -0.83 | omega-hydroxypalmitate O-feruloyl transferase |
| Medtr6g086365 | -1.16 | n.r. | -2.20 | n.r. | arabinogalactan protein |
| Medtr6g086390 | n.r. | n.r. | -1.59 | n.r. | arabinogalactan peptide-like protein |
| Medtr4g059720 | n.r. | n.r. | n.r. | 0.83 | fasciclin-like arabinogalactan protein |
| Medtr5g081810 | n.r. | n.r. | n.r. | 1.18 | laccase/diphenol oxidase family protein |

Firstly, we focused on the regulation of Strigolactone (SL) signalling. SL are carotenoid-derived plant hormones (Gomez-Roldan et al., 2008; Umehara et al., 2008) that also function as extraradical signals to activate AM fungi (Waters et al., 2017). SL biosynthesis (*CCD8-1* and *CCD8-2*) and transport (*PDR1a*) marker genes were down-regulated at 28 dpi in MYC compared to CTR plants (Fig. 2a), which is in line with the inhibition of rhizospheric SL signaling upon maximum root colonization (Lopez-Raez et al., 2011). Remarkably, this down-regulation of SL biosynthesis and transport genes occurred as early as 10-14 dpi in CO-treated plants (MYC+CO *vs* CTR). A rather different scenario was recorded in CO-treated plants that were not inoculated with AMF (CTR+CO *vs* CTR, SI_DATA1). Under these conditions, SL biosynthesis and transport related genes were found to be up-regulated between 14 and 21 dpi. Such an up-regulation of SL biosynthetic genes suggests a CO-dependent boosting of fungus-directed signaling in the absence of an effective interaction with the symbiont.

**Fig. 2.**
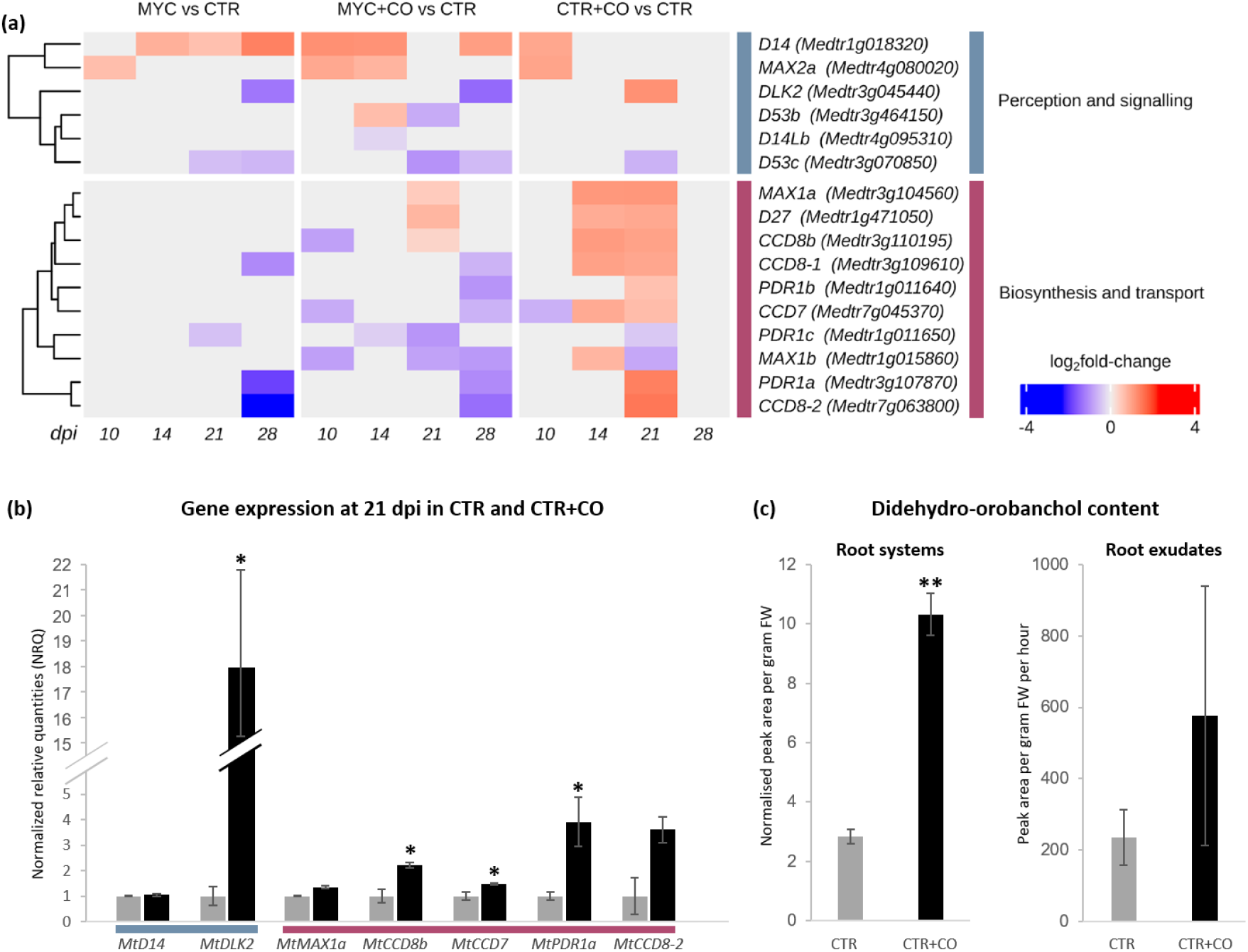
CO effect on SL-related gene expression and SL accumulation. **(a)** Heatmap plot of strigolactone-related genes in each comparison and time points. A progressive decrease in strigolactone biosynthesis (blue) is in line with the progress of AM development in MYC *vs* CTR. This down-regulation was earlier and more extensive in MYC+CO *vs* CTR comparison, in line with the acceleration of colonization observed in the presence of CO treatment. Remarkably, a strong up-regulation (red) was evident in both strigolactone biosynthesis and transport until 21 dpi in the CTR+CO *vs* CO comparison. Genes are clustered within each category according to Spearman’s distance (Müller et al., 2019). **(b)** Independent experiment confirming SL gene expression analysis in CO treated and untreated roots at 21 dpi. Mean values ± SEs of three biological replicates for each treatment are shown. Asterisks indicate statistically significant differences (Student’s t-test, *P < 0.05). **(c)** Didehydro-orobanchol content in root tissues and exudates of CTR and CTR+CO plants sampled at 21 dpi. A significant increase in DDO root content correlated with CO treatment. A similar trend was observed in the exudates, even if the difference between CTR and CTR+CO plants was statistically not significant. Mean values ± SDs of four biological replications for each treatment are shown. Each biological replication was composed by two plants. Asterisks indicate statistically significant differences (Student’s t-test, **P < 0.01).

An independent experiment confirmed the same pattern of gene regulation at 21 dpi in CTR+CO compared to CTR plants, with an unchanged expression of *MtD14*, a significant upregulation of *MtDLK2, MtCCD8b, MtCCD7* and *MtPDR1a* and a comparable (albeit not significant) trend for *MtMAX1a* and *MtCCD8-2* (Fig. 2b).

To correlate the observed regulation of SL related gene expression to SL metabolomics, we analyzed SL content in root tissues and exudates from 21-day-old CTR and CTR+CO plants. In our samples only one SL, didehydro-orobanchol (DDO), was above the detection level. This SL has been identified in roots of several AM host plants (Lopez-Raez et al., 2008; Yoneyama et al., 2008; Kohlen et al., 2012), and characterized as the major strigolactone in *M. truncatula*, with a role in stimulating hyphal branching in the AM fungus *Gigaspora margarita* (Liu et al., 2011, Tokunaga et al., 2015). Interestingly, CTR+CO roots contained a significantly higher level of DDO compared to CTR roots. A comparable trend (albeit statistically not significant) was also observed in root exudates (Fig. 2c).

Secondly, MYC+CO *vs* CTR and MYC+CO *vs* MYC comparisons (Table 1; SI_DATA4, 5) highlighted the up-regulation of a consistent group of functional categories related to cell remodelling, such as *Cytoskeleton Organization*, and *Cell Wall Organization*. This is in line with the activation of intracellular fungal accommodation by the host tissues (Luginbuehl and Oldroyd, 2017; Pimprikar and Gutjarh, 2018), requiring the reorganization of cytoskeletal elements (Genre et al., 2005), the onset of massive exocytic processes (Genre et al., 2012), cell wall material deposition and remodelling (Balestrini and Bonfante, 2014) to generate the symbiotic interface (Balestrini et al., 2005). Furthermore, a significant up-regulation was also observed in *Cell Cycle Organization* for numerous cell cycle regulators (*CYCA*, *CYCB*, *alpha-AURORA kinase activator*) and cell plate-associated proteins (*KNOLLE*), in strong agreement with the described activation of cell division (Russo et al., 2018) and endoreduplication (Carotenuto et al., 2019) as part of the fungal accommodation response.

In the light of this data, and because fungal accommodation responses such as prepenetration apparatus (PPA) development are known to occur in the range of a few hours (Genre et al., 2005), we decided to further investigate earlier responses to CO, by applying local, 6 hour long treatments to *M. truncatula* ROCs (Fig. 3) and analyzing the regulation of *MtKNOLLE* (Richter, et al., 2014; Russo et al., 2019) and two additional cell cycle-related markers known to be expressed in early AM development, *MtAPC2*, the *alpha subunit of the Adaptor Protein complex2* (Van Damme et al., 2011; Russo et al., 2018) and *MtCYCL3*, a cyclin-like F-box protein (Russo et al, 2019). Two additional early symbiotic markers were also analyzed: *MtPUB1* (*Plant U-box protein1*), an E3 ubiquitin ligase (Vernié et al., 2016) and *MtCBF3*, a CAAT box-binding transcription factor (Hogekamp et al., 2011).

**Fig. 3.**
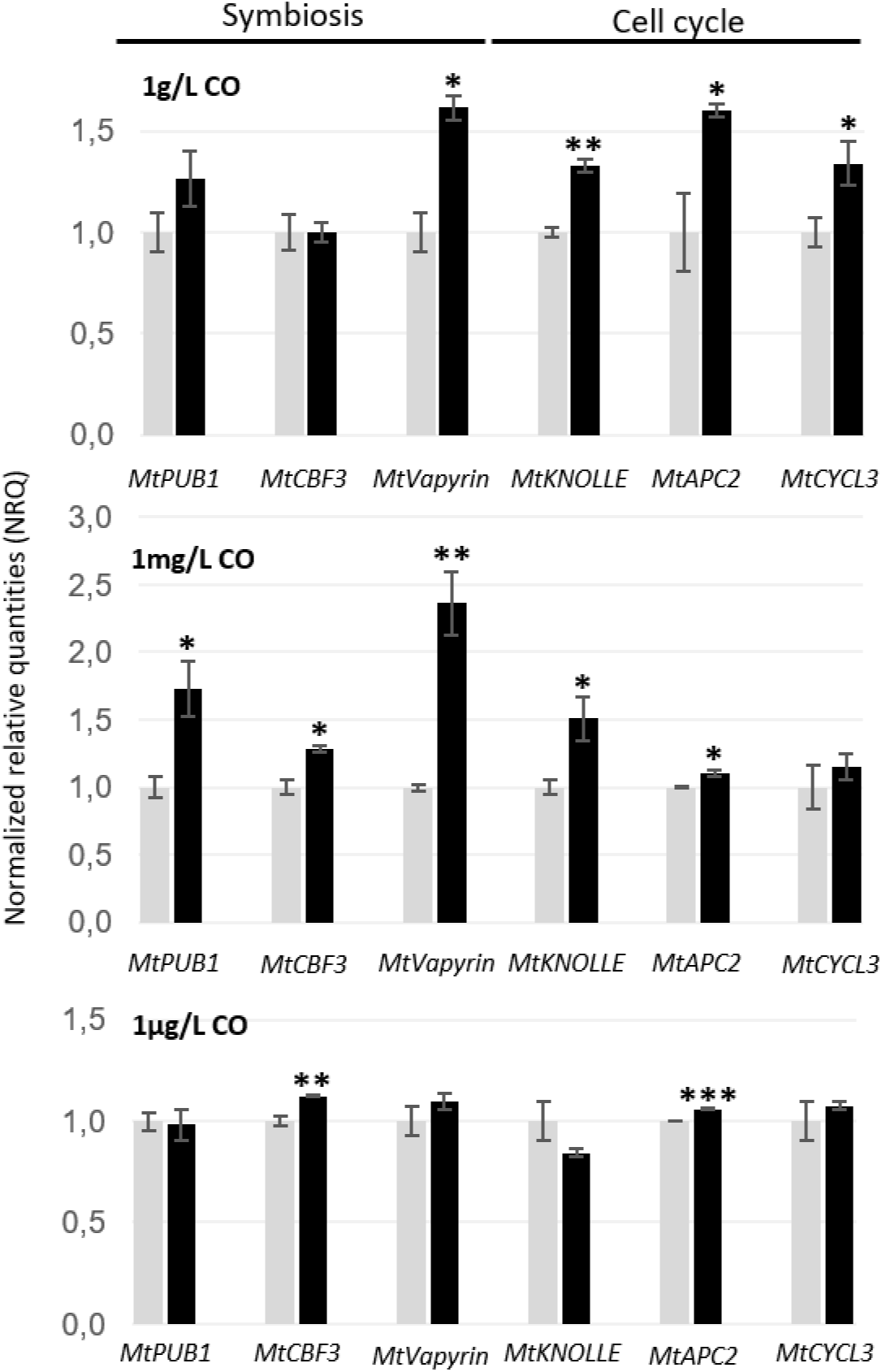
Local regulation of early AM marker genes. Local CO treatment stimulated concentration-independent expression of several genes involved in fungal accommodation. The highest number of significantly upregulated genes was recorded upon 1mg/L CO application. Grey bars = control, water treated roots; black bars = CO treated roots. Mean values ± SEs of five biological replicates for each treatment are shown. Asterisks indicate statistically significant differences (Student’s t-test, *P < 0.05, **P < 0.01, and ***P < 0.001).

As shown in Figure 3, each gene was upregulated upon either 1 g/L, 1 mg/L or 1 μg/L CO treatment, with 1 mg/L CO solution resulting to significantly up-regulate *MtPUB1, MtCBF3, MtVapyrin, MtKNOLLE* and *MtAPC2*, whereas *MtCYCL3* was significantly induced only in response to 1g/L treatment. By comparing these results with the available data on the Noble *MtGEA V3* database (Bendito et al., 2008; https://lipm-browsers.toulouse.inra.fr/pub/expressionAtlas/app/mtgeav3), we observed a general overlap with the regulation of the same genes during early root-fungus contact (Ortu et al., 2012) and 6 h LCO treatment (Czaja et al., 2012).

In conclusion, our targeted investigation of gene expression confirmed the CO-dependent activation of several early AM markers and revealed that a 6 h CO application is sufficient to activate a set of genes related to prepenetration responses. Together, this gene expression pattern corroborated our RNA-seq data and was fully compatible with the observed acceleration of AM development in CO-treated plants, providing molecular evidence in favor of a CO-dependent advance of pre-penetration responses in root cells.

### CO treatment stimulated pre-penetration responses upon AM inoculation

We then decided to investigate the effect of CO treatment on prepenetration responses in the presence of an AM fungal inoculum. To this aim, we grew *M. truncatula* ROCs expressing GFP-HDEL in the presence of 1 mg/L CO solution (the most active concentration in our gene expression analyses) or sterile water as a control. GFP-HDEL is a fluorescent marker for the endoplasmic reticulum (ER) that has previously been used to track cytoplasmic aggregations and PPA development during AM colonization (Genre et al., 2005; 2008; 2012). The asynchronous progression of hyphal development and root colonization limits our ability to fully control the timing of fungus-plant contact even using the so-called targeted inoculation method that we here adapted (Chabaud et al., 2002). For this reason, observations were done at 7, 10 or 14 dpi with pre-germinated *G. margarita* spores, a time frame that corresponds to hyphopodium formation and initial epidermis colonization. Using hyphopodia as a hallmark to locate plant-fungus contact sites, we focused our observations on the surrounding epidermal cells. As expected, broad ER aggregations were observed at all time points, extending between the hyphopodium contact site and the epidermal cell nucleus, as previously described during PPA formation (Genre et al., 2005). Such aggregations were observed both in the presence or absence of the CO treatment. Nevertheless, such PPA-related ER aggregations appeared to be more frequent in CO-treated roots, as confirmed by quantitative analysis (Fig. 4). Indeed, the percent of cells showing ER aggregations in hyphopodium-contacted areas was significantly higher in treated than untreated roots both at 7 and 10 dpi, while the two data were comparable at 14 dpi. The major impact of CO treatment on earlier time points can be explained with the progressive decrease in residual CO concentration and/or the inhibition of new hyphopodium formation as arbuscules start developing in the inner root tissues. In this scenario, the observed significant stimulation of prepenetration responses in early time points provides a functional validation of our transcriptomic data (Table 1) and the first cellular basis for the observed CO-dependent promotion of AM colonization (Volpe et al., 2020).

**Fig. 4.**
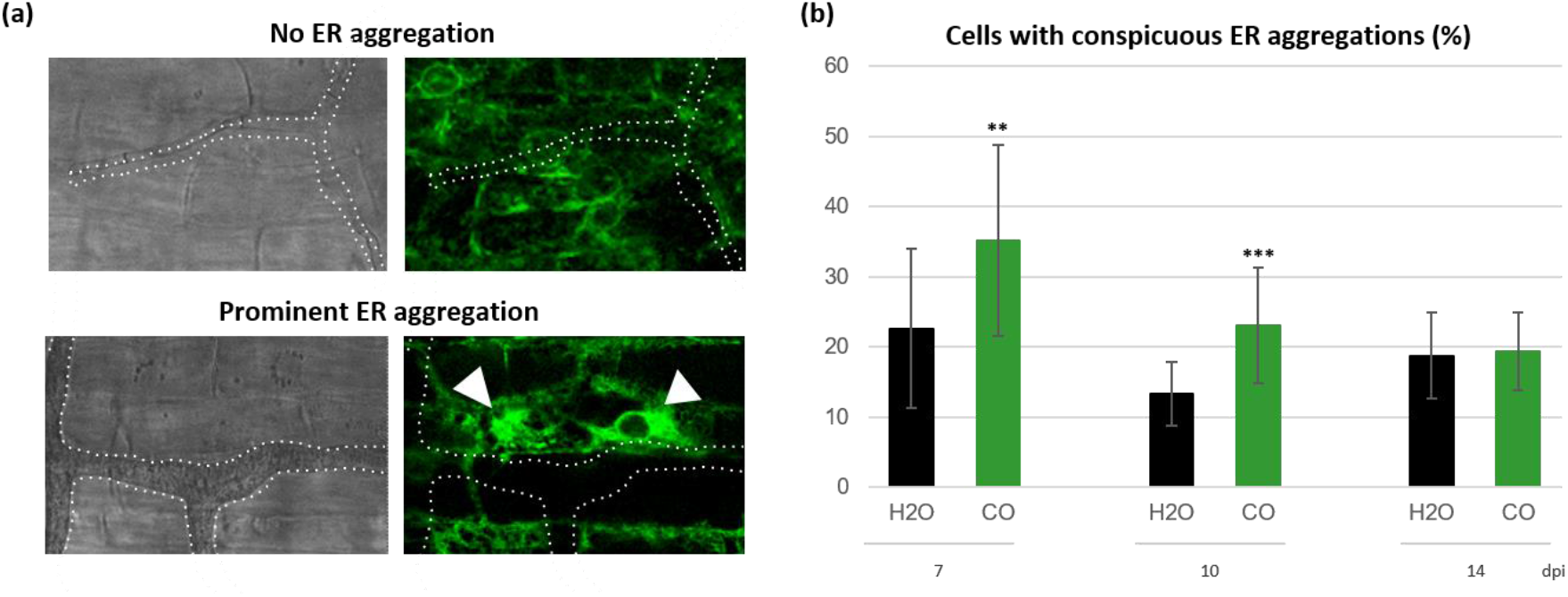
PPA formation in epidermal cells of *M. truncatula*. Root organ cultures expressing GFP-HDEL in the endoplasmic reticulum (ER) were treated with 1mg/L CO or water (H_2_O) as control at the time of inoculation with pre-germinated *G. margarita* spores. Epidermal cells in the vicinity of fungal hyphopodia were then imaged at 7, 10, 14 dpi and categorized based on the absence or presence of ER aggregations **(a)**. A statistically significant increase in the percentage of epidermal cells with prominent ER aggregation was recorded in CO-treated roots at 7 and 10 dpi **(b)**. At least 100 independent infection events were visualized for each treatment. Asterisks indicate statistically significant differences (Student’s t-test, *P < 0.05, **P < 0.01, and ***P < 0.001).

### Long term effects of the CO treatment

The 28 dpi time point was associated with a significant increase in plant development for MYC+CO compared to MYC plants (Fig. S7a). According to our previous studies (Volpe et al., 2020), this is also the time of maximum AM development in our pot cultured *M. truncatula*. This was confirmed by our quantitative analysis, which also highlighted a significant promotion of AM colonization in MYC+CO compared to MYC plants (Fig. S7b). Nevertheless, when comparing MYC+CO to MYC root transcriptome and to MtExpress V3 gene expression data (Carrere et al., 2021), we observed a surprising decline in the expression of several AM markers, including transcription factors (*MtMYB* and *MtRAM2*) and arbuscule-specific phosphate (*MtPT4*) and ammonium (*MtAMT1*) transporters (Fig. 5a; SI_DATA4). We suspected that this could indicate the inception of symbiosis senescence, possibly related to space limitations to root development in our pot cultures: in fact, the root system of MYC+CO plants had extended to the whole substrate volume at 28 dpi, which was not the case for MYC plants.

**Fig. 5.**
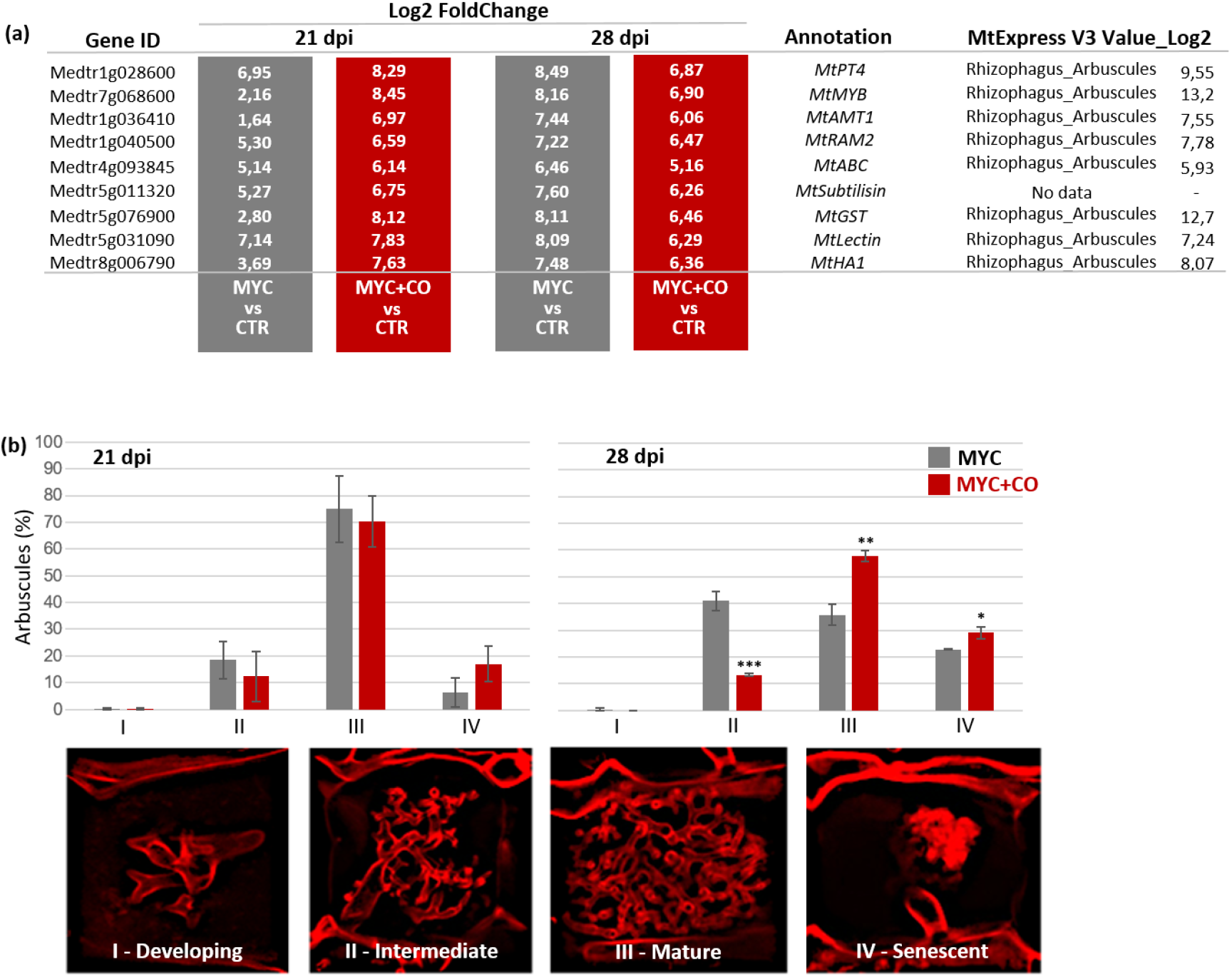
Quantitative analyses of mycorrhizal phenotypes between MYC and MYC+CO plants. **(a)** Expression of AM-induced genes increased at 21 but declined at 28 dpi in the comparison MYC+COvsCTR compared to MYCvsCTR, suggesting the onset of arbuscule senescence at the latest time point. Normalized log2 expression values from Expression Atlas of Medicago (MtExpress V3; Reference dataset 20220901; https://lipm-browsers.toulouse.inra.fr/ pub/expressionAtlas/app/v3/) are reported in the right column **(b)** Arbuscules were imaged at 21 and 28 dpi in MYC and MYC+CO plants and classified based on their developmental stage: developing arbuscules with a trunk and very few branches (stage I), intermediate arbuscules with thick branches (stage II), fully mature arbuscules with abundant fine branches (stage III) and senescent arbuscules with clusters of collapsed branches (stage IV). A comparable distribution between MYC and MYC+CO plants was observed at 21 dpi, even if a non-significant reduction and increase were respectively observed in Stage II and Stage IV arbuscules. Such trends became statistically significant at 28 dpi, associated with an increase in the abundance of Stage III arbuscules in MYC+CO plants. Data are based on the measurements of 30 infection units for each of four biological replicates. Mean values ± SDs of four biological replications for each treatment are shown. Asterisks indicate statistically significant differences (Student’s t-test, *P < 0.05, **P < 0.01, and ***P < 0.001).

We therefore used confocal microscopy to image and compare arbuscule morphology in MYC and MYC+CO root samples. To obtain a more dynamic view of the colonization process, we extended our analysis to both 21 and 28 dpi. Selected roots were stained with WGA-FITC for detailed imaging of the fungal cell wall and individual arbuscules were classified into four developmental stages based on their morphological features to generate a quantitative comparison (Fig. 5b): Stage I (developing arbuscules with a limited number of large branches); Stage II (intermediate maturity, with several fine branching occupying part of the host cell lumen); Stage III (mature arbuscules, with fine branches in most of the host cell volume); Stage IV (senescent arbuscules displaying clusters of collapsed branches). As shown in the histogram of Figure 5b, arbuscule morphology in MYC and MYC+CO plants was comparable at 21 dpi, even if a non-significant decrease in Stage II and an increase in Stage IV arbuscules was observed. Interestingly, this trend became more evident - and statistically significant - at 28 dpi, where the arbuscule population in MYC+CO plants also displayed a significant increase in Stage III. At 28 dpi, this resulted in a shift in the distribution of arbuscule classes, with a maximum frequency in Stage II for MYC plants and in Stage III for MYC+CO plants. Altoghether, we interpret this developmental shift as the consequence of the CO-induced advancement in fungal accommodation responses - as indicated by our molecular analyses and further demonstrated in epidermal cells by live cell imaging - culminating in a significant advance in arbuscule development and senescence in CO-treated plants.

### CO promotion of AM development depends on DMI3

In a previous study we demonstrated the promotion of AM colonization by CO treatment (Volpe et al., 2020), confirmed also in this study (Fig. S7b). In order to verify whether the observed effect was dependent on CSSP-mediated CO perception, we now applied the same CO treatment to *dmi3-1* mutant plants (Lévy et al, 2004; Mitra et al, 2004) and compared the AM phenotype at 28 dpi between treated and untreated roots. The *M. truncatula Dmi3* gene encodes a nuclear localized Ca^2+^ and calmodulin-dependent kinase that is a central hub of the CSSP (Choi et al., 2018). Its mutation blocks AM colonization at the root epidermis. As described in literature, the only recognizable fungal structures associated with *dmi3-1* roots were hyphopodia, while no intraradical fungal development was observed in any of our samples, independent of the CO treatment. Furthermore our quantitative analysis revealed a comparable abundance of hyphopodia developed in the presence or absence of the CO treatment (Fig. S8a). Such a lack of any evident effect associated with the CO treatment in the *dmi3-1* mutant, strongly suggested that the effects observed in WT plants depend on the CSSP activity.

In an additional set of experiments, we investigated the transcriptional response to COs in *dmi3-1* ROCs, by testing the expression of early AM markers (*MtPUB1, MtCBF3, MtVapyrin*) and prepenetration-related genes (*MtKNOLLE*, *MtAPC2*, *MtCYCL3*; Fig. S8b). The regulation of five out of those six genes (*MtCBF3, MtVapyrin, MtKNOLLE, MtAPC2* and *MtCYCL3*) was comparable to that reported in the central panel of Fig. 3 for the WT. This upregulation of AM responsive genes in *dmi3* mutants exposed to AM fungal signals has indeed been reported in *M. truncatula* (Kosuta et al., 2003) and rice (Gutjahr et al., 2008), and interpreted as clue to the existence of alternative signaling, acting in parallel to at least one part of the CSSP. A remarkable exception is represented by *MtPUB1*, which was significantly induced in WT samples but downregulated in *dmi3-1* mutants, in line with the canonical CSSP-based model of AM signaling. This pattern of gene regulation, including *Dmi3*-dependence of *MtPUB1*, is comparable to the results of previous studies on early root-fungus contact (Ortu et al., 2012) and 6 h LCO application (Czaja et al., 2012).

## Discussion

The main aim of this investigation was to test the hypothesis that CO stimulate symbiotic responses in the host root throughout AM development. In fact, we recently demonstrated that early CO treatment promotes root infection by AM fungi over 28-48 days (Volpe et al., 2020), but a major gap was left in our understanding of the impact of CO treatment on root transcriptome, metabolism and cellular responses throughout AM colonization.

Indeed, several studies have investigated early plant responses to exogenous treatment with AM fungal raw exudates or purified LCO and CO, revealing the rapid (< 1h) activation of plant symbiotic signaling processes (Maillet et al., 2011; Genre et al., 2013) and short-term (1-48 hours) transcriptional reprogramming (Czaja et al., 2012; Giovannetti et al., 2015; Feng et al., 2019). All of these investigations were anyway analyzing plant responses occurring in the absence of any fungal inoculation.

By analyzing the root transcriptome alongside symbiosis development in a period of over four weeks following initial CO treatment, the present study breaks through this limitation. The consolidation of our gene expression analysis with functional insights generated a first, consistent outlook on CO-dependent plant responses. At any rate, our large transcriptomic dataset, combining four different conditions and four time points, remains available (link in Data Availability Statement) to drive further investigations.

### Exogenous CO stimulate plant symbiotic responses throughout AM development

Our present results demonstrate that, besides the well-characterized activation of early symbiotic signaling and gene regulation, CO also influenced key symbiotic features of the host tissues from as soon as 6 hours after treatment to 28 dpi. Altogether, CO impacted on the expression of root genes related to SL metabolism and transport, altered root SL content; repressed pathogenesis-related genes and actors of effector-triggered immunity and gene pathways involved in secondary metabolism (isoflavonoid and terpenoid synthesis); increased the expression of lipid and mineral nutrient transporters, globally suggesting the reinforcement of the host response to the AM fungus and the promotion of a symbiosis-oriented metabolic and physiological context. Furthermore, CO also promoted the expression of genes involved in cell-cycle reactivation and fungus accommodation, in association with an observed induction of prepenetration responses, and caused an advance in the root colonization process that extended to 28 dpi.

In short, our experimental results indicate a general and long-lasting stimulation of symbiotic responses in both inoculated and uninoculated CO-treated plants, demonstrating that CO have all the predicted features of Myc-factors, intended as fungal-secreted molecules that prepare the host plant to symbiosis establishment (Gutjahr & Parniske 2013) and actively promote it.

### The strange case of an unattainable love

Root exposure to fungal signaling molecules in the absence of the fungus (our CTR+CO condition) represents a markedly artificial situation. In nature, as AM fungal hyphae approach a host root, Myc-factor perception takes place alongside (or shortly before) additional, largely uncharacterized signals (Bonfante and Requena, 2011), including the physical contact between hyphae and root epidermal cells, and the subsequent development of intraradical colonization (Choi et al., 2018).

The progression of AM development has an impact on root responses, by transiently triggering defense reactions (Garcia-Garrido and Ocampo, 2002; Jung et al., 2012; Cameron et al., 2013), progressively reducing SL synthesis and secretion (Lopez-Raez et al., 2011; this paper) and overall outlining the context of a successful symbiotic interaction. By contrast, our CTR+CO plants lack this second set of fungal stimuli, which are crucial modulators of the host response. When interpreting the gene regulation pattern in these root samples, the presence of a certain degree of overreaction should therefore be taken into account. With this caveat, the CTR+CO condition provided the experimental tool to dissect otherwise unseen CO effects. A remarkable example of this artificial amplification of plant responses is the boosting of SL-related gene expression, which extended for a few weeks after CO treatment in CTR+CO plants. We interpret this as a climax in plant presymbiotic signaling, in a condition where physical plant-fungus interaction is not going to occur. While the biological significance of this observation can be limited, with reference to a natural context, it convincingly indicates that CO effects extend over several weeks and impact on SL content, a root feature that is directly linked to AM symbiosis establishment.

### A symbiotic signaling loop

Plants exude diverse mixtures of SL, depending on plant species, growth conditions and developmental stages (Wang and Bouwmeester, 2018). With reference to AM signaling, DDO has been characterized as the most active molecule in *M. truncatula* (Lopez-Raez et al., 2008; Yoneyama et al., 2008; Kohlen et al., 2012). In this frame, our novel observation of SL metabolism-related gene upregulation upon CO treatment is further reinforced by DDO accumulation in the root, suggesting a redirection of SL metabolism towards AM fungus-directed signals in CO-treated roots. Furthermore, previous investigations have recorded an increase of CO release by SL-treated AM fungi (Genre et al., 2013). Altogether, this suggests the existence of a positive feedback mechanism that reinforces reciprocal plant-fungus signaling in support of symbiosis establishment.

To summarize our conclusions, we propose a model - schematized in Figure 6 - where the exogenous application of CO does not only boost signal exchange in this symbiotic signaling loop, but also stimulates intracellular accommodation responses in the host root, facilitating fungal colonization and eventually accelerating symbiosis development.

**Fig. 6.**
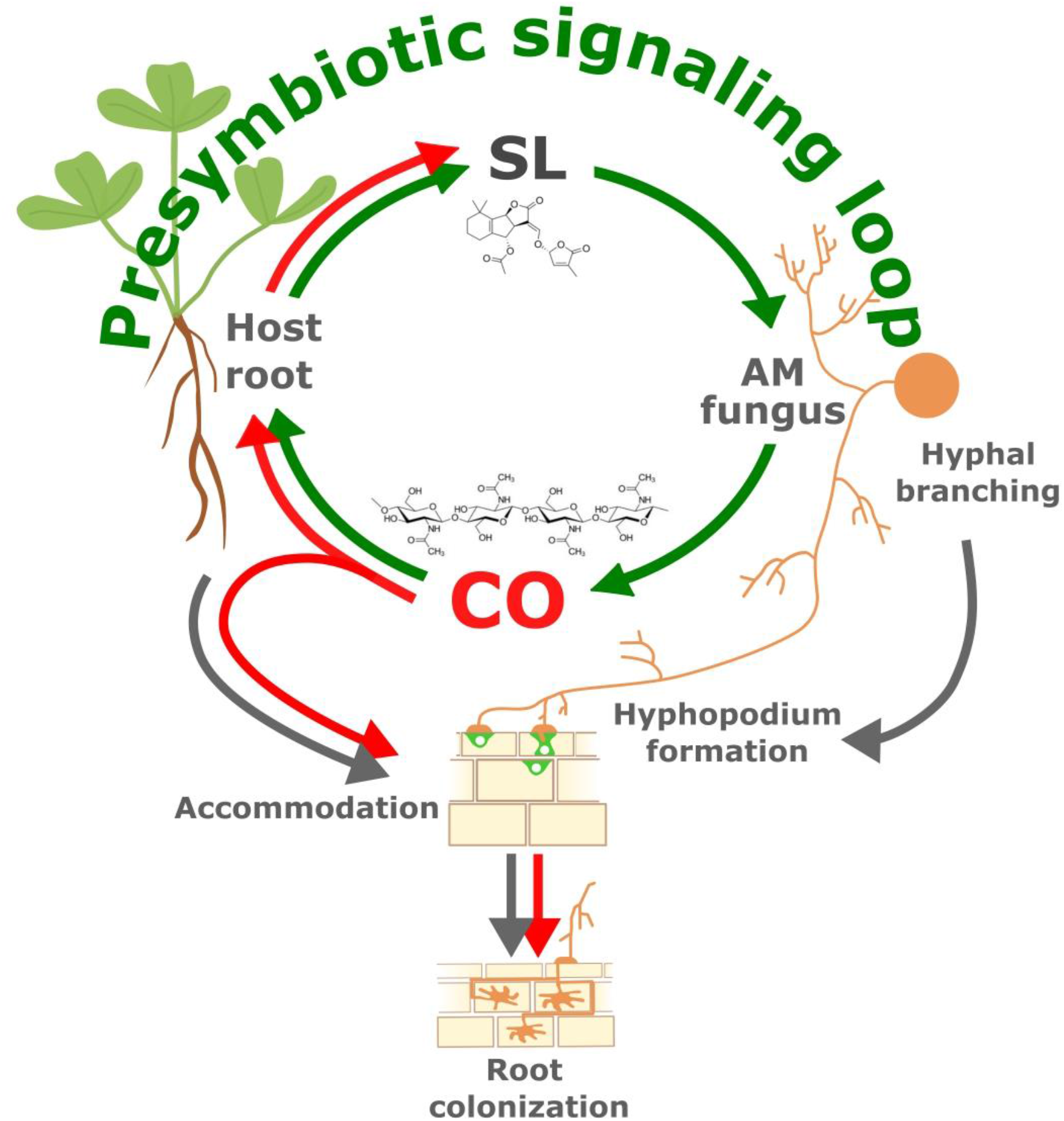
Schematic representation of exogenous CO effects on AM symbiosis development. The perception of root-released SL by AM fungi is known to induce hyphal branching and boost CO release. In turn, CO perception activates root symbiotic responses leading to intracellular fungal accommodation and symbiosis establishment. Our current results (red arrows) indicate that exogenous CO application further stimulates SL biosynthetic gene expression and SL accumulation in the host root, suggesting the presence of a positive feedback mechanism that we propose here as a presymbiotic signaling loop. As an additional consequence of CO application, we also observed a stimulation of intracellular fungal accommodation, supporting the observed advance in root colonization and symbiosis development.

## Acknowledgments

We are grateful to David Barker and Paola Bonfante for constructive discussion and their critical revision of the manuscript draft. We also thank David Barker to kindly providing *Medicago dmi3-1* mutant seeds. This research was funded by Fondazione Cassa di Risparmio di Cuneo (Bando Ricerca Scientifica 2015–Project AMforQuality 2015/17271). We are grateful to Erik Limpens for hosting LC in his lab in Wageningen for an Erasmus+ traineeship.

## Author Contributions

VV, TM performed experiments and data analysis. MC performed data analysis and contributed figure editing and manuscript writing. AC, SC, LC and WK performed experiments. VV, AG conceived experiments and wrote the manuscript.

## Data Availability

Raw RNA-seq data were deposited in the Sequence Read Archive (SRA, NCBI) under accession PRJNA813377. Data will be released upon manuscript acceptance and are temporarily available to reviewers at the following link: https://dataview.ncbi.nlm.nih.gov/object/PRJNA813377?reviewer=h8u5q75tpu8t65mqi52nk90h52

