## Supporting Information for "Long-lasting impact of chito-oligosaccharide application on strigolactone biosynthesis and fungal accommodation promotes arbuscular mycorrhiza in *Medicago truncatula*"

The following Supporting Information is available for this article:

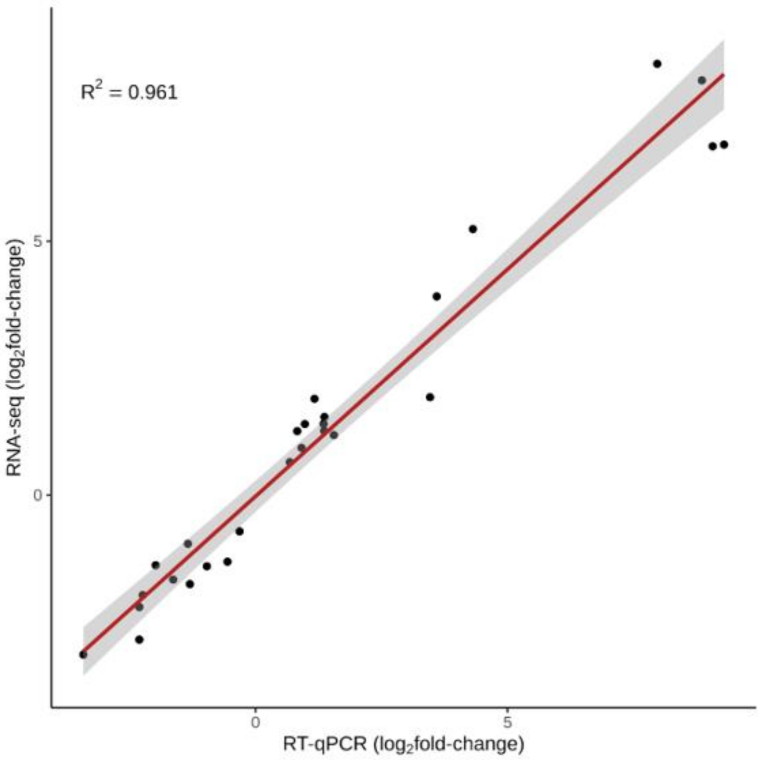


**Fig. S1 RNAseq and qRTPCR data correlation.** Values of $log_2$fold-change for 10 differentially expressed genes measured by qRT-PCR correlate with fold-change values emerged in the RNA-seq analysis ($R^2$ = 0.961). Differentially expressed genes (both up- and down-regulated) were randomly chosen (see materials and methods) across all comparisons (CTR+CO *vs* CTR, MYC *vs* CTR, MYC+CO *vs* CTR, MYC+CO *vs* MYC) and time points (10, 14, 21, 28 dpi).

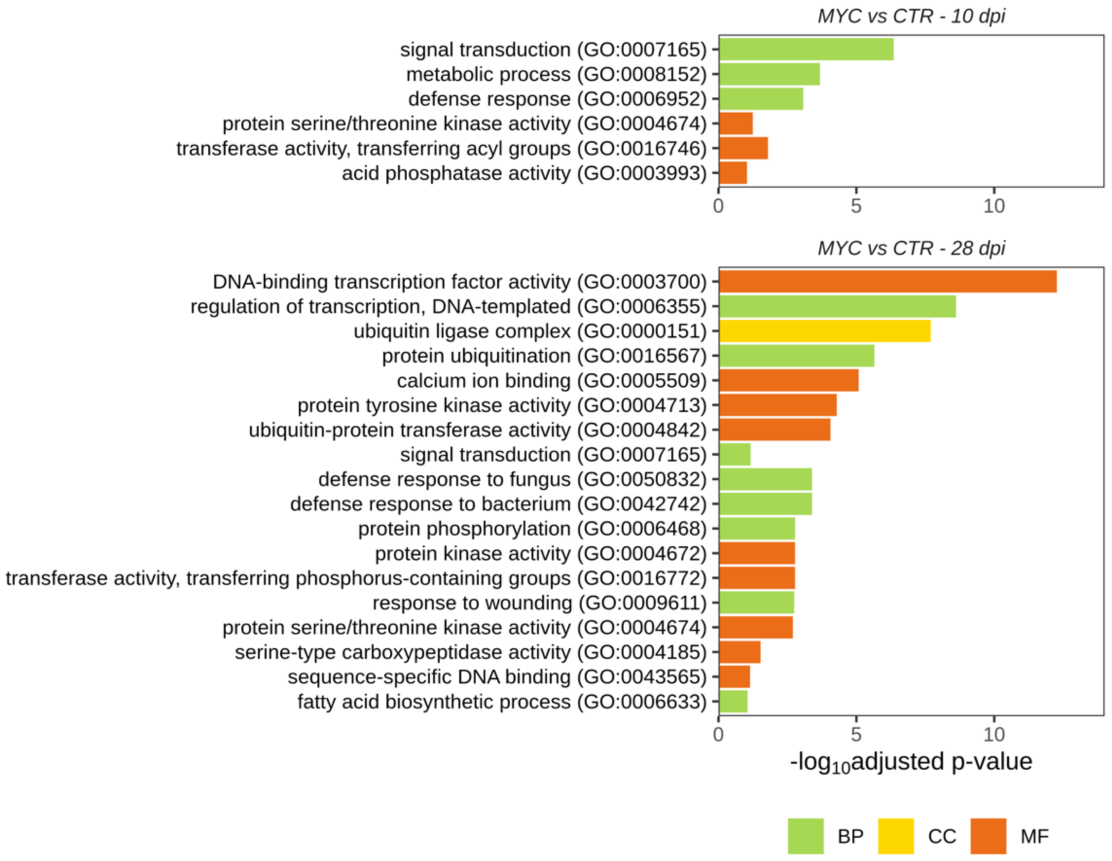


**Fig. S2 Gene Ontology (GO) enrichment analysis in the comparison MYC *vs* CTR at 10 and 28 dpi.** Green = biological process (BP); Yellow = cellular component (CC); Orange = molecular function (MF).

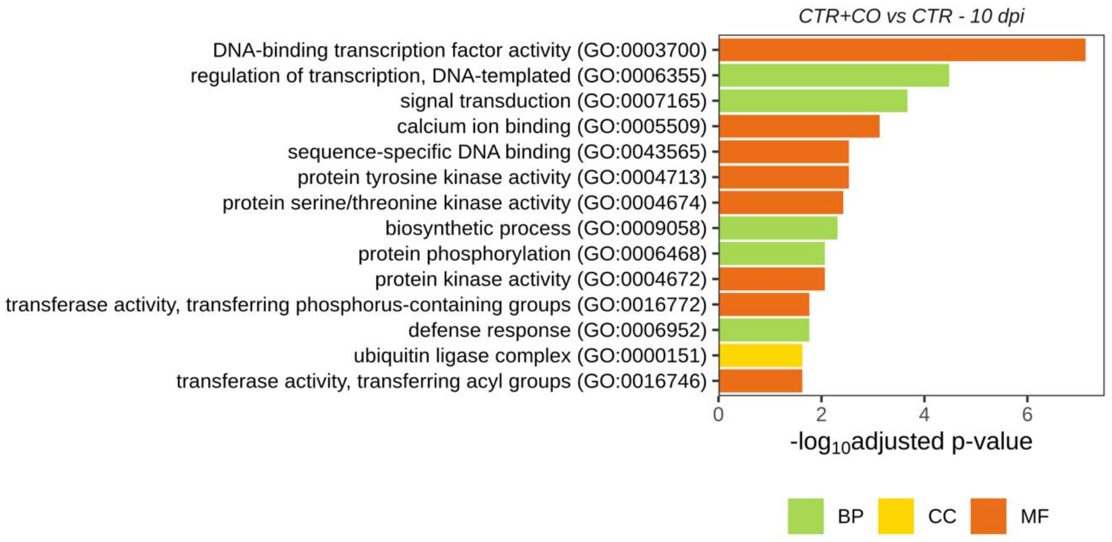


**Fig. S3 Gene Ontology (GO) enrichment analysis in the comparison CTR+CO *vs* CTR at 10 dpi.** Green = biological process (BP); Yellow = cellular component (CC); Orange = molecular function (MF).

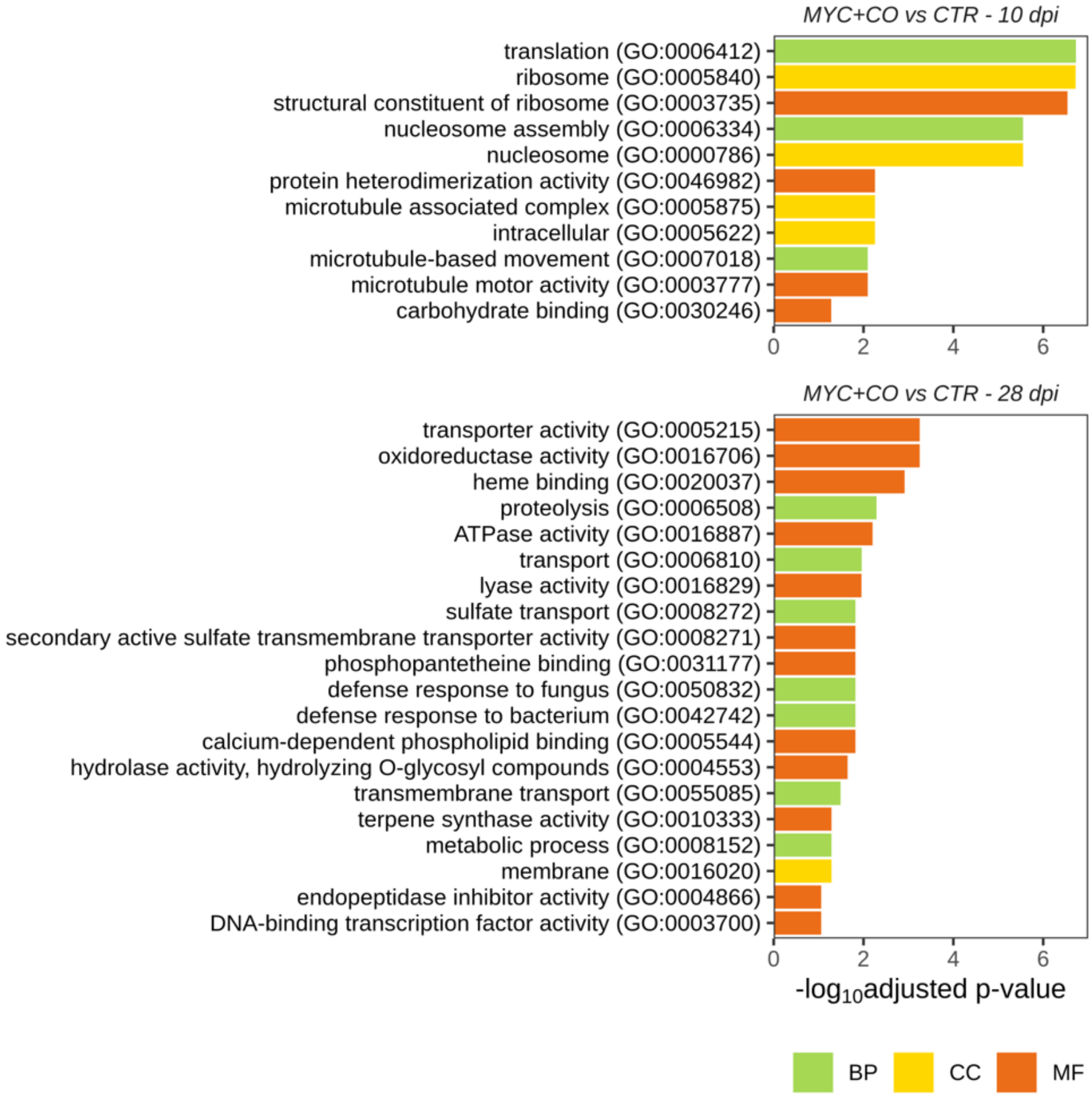


**Fig. S4 Gene Ontology (GO) enrichment analysis in the comparison MYC+CO *vs* CTR at 10 and 28 dpi.** Green = biological process (BP); Yellow = cellular component (CC); Orange = molecular function (MF).

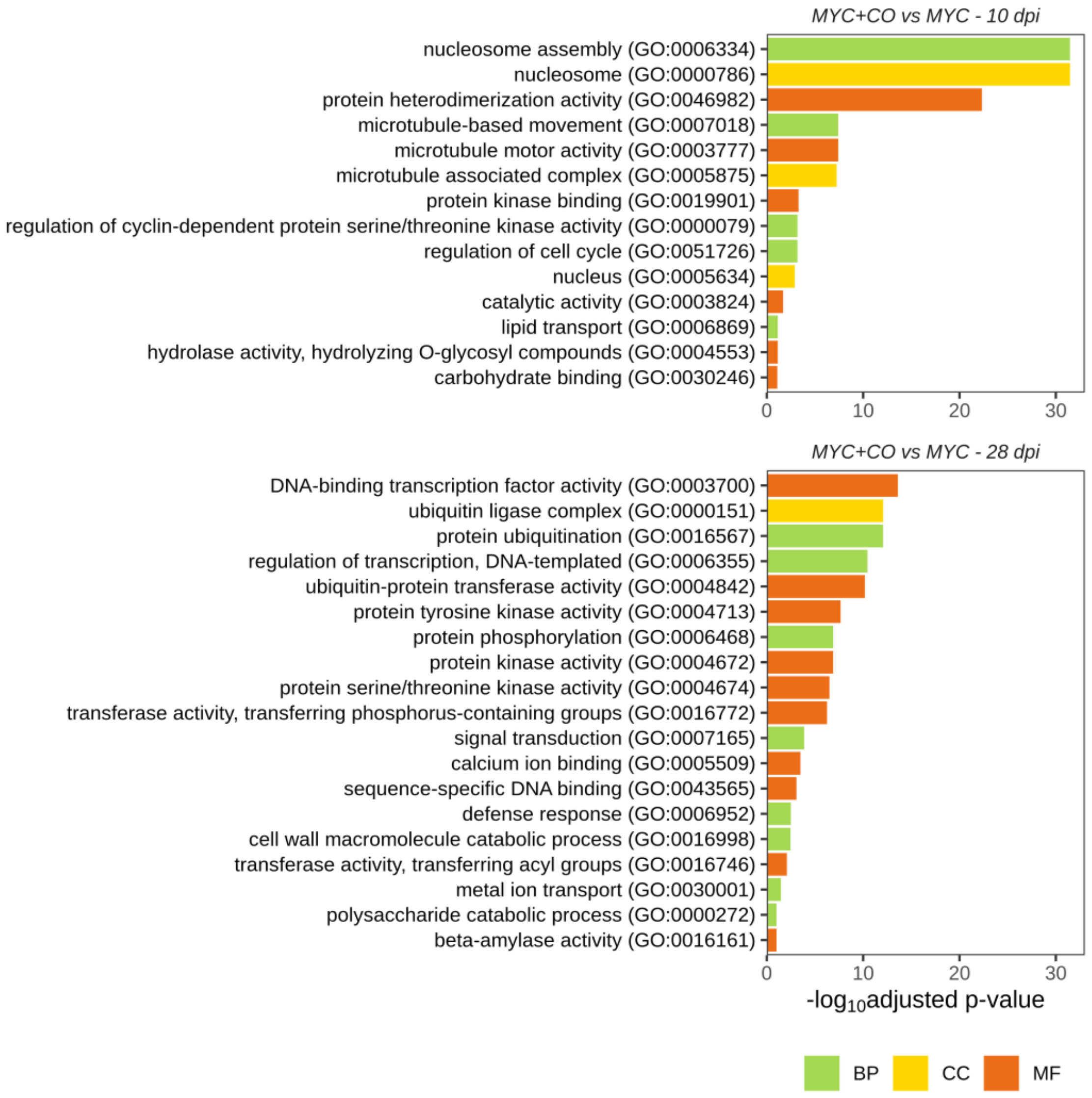


**Fig. S5 Gene Ontology (GO) enrichment analysis in the comparison MYC+CO *vs* MYC at 10 and 28 dpi.** Green = biological process (BP); Yellow = cellular component (CC); Orange = molecular function (MF).

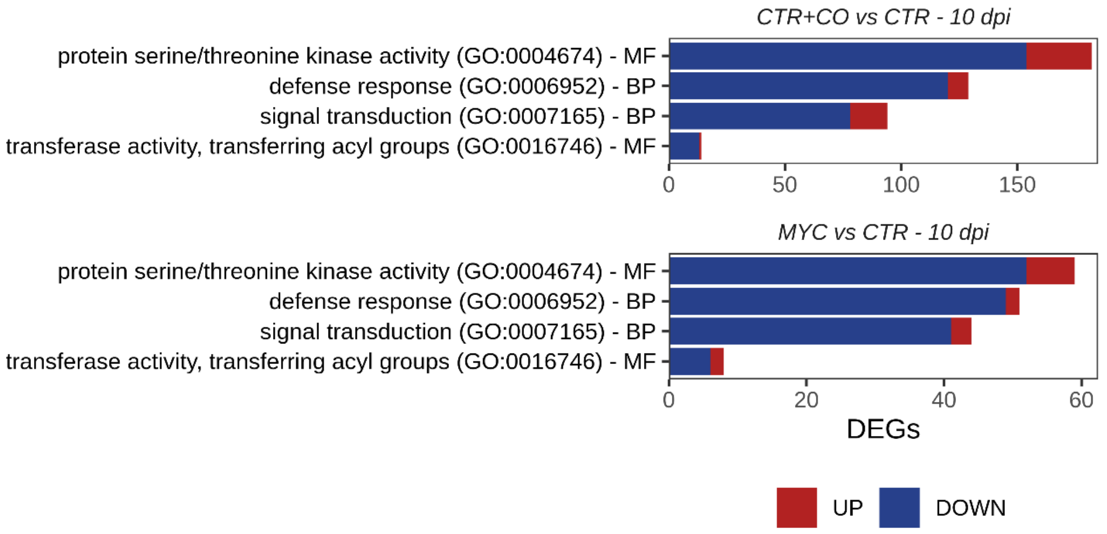


**Fig. S6 Gene regulation pattern in CTR+CO and MYC plants compared to CTR.** GO enrichment analysis revealed that the global regulatory pattern of four among the most represented functional categories at 10 dpi displays comparable trends in response to either AM inoculation or CO treatment. Red = upregulated genes; blue = downregulated genes.

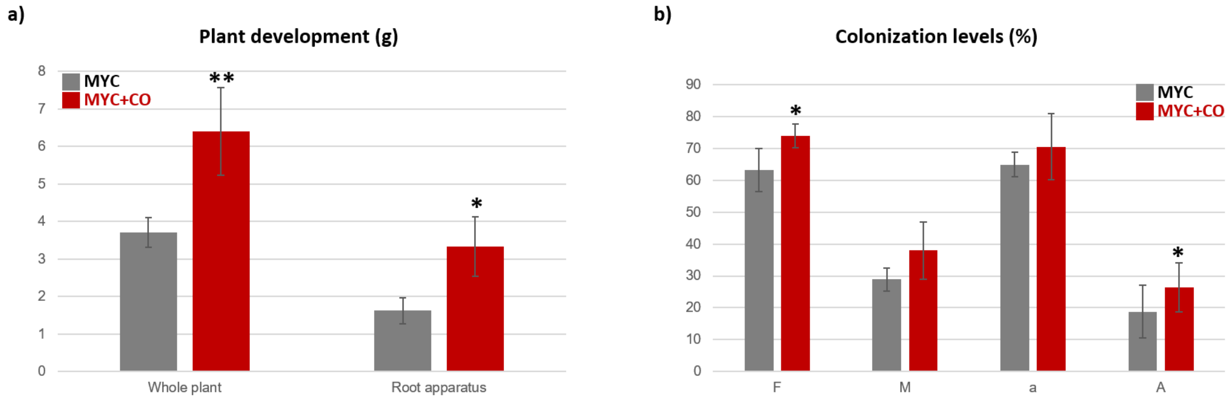


**Fig. S7 Morphological analysis in mychorrized plants.** a) Plant fresh biomass analysis. A significant increase in whole plant and root apparatus development was recorded in MYC+CO compared to MYC plants. Error bars indicate standard deviation. Asterisks indicate statistically significant differences (Student's t-test, *p<0,05; **p<0,005). b) Colonization levels in MYC and MYC+CO plants. A significant increase in terms of frequency (F) and arbuscule abundance (A) was observed in MYC+CO plants compared to MYC ones. Histograms represent average values for each parameter; error bars indicate standard deviation (Student's t-test, *p<0,05).

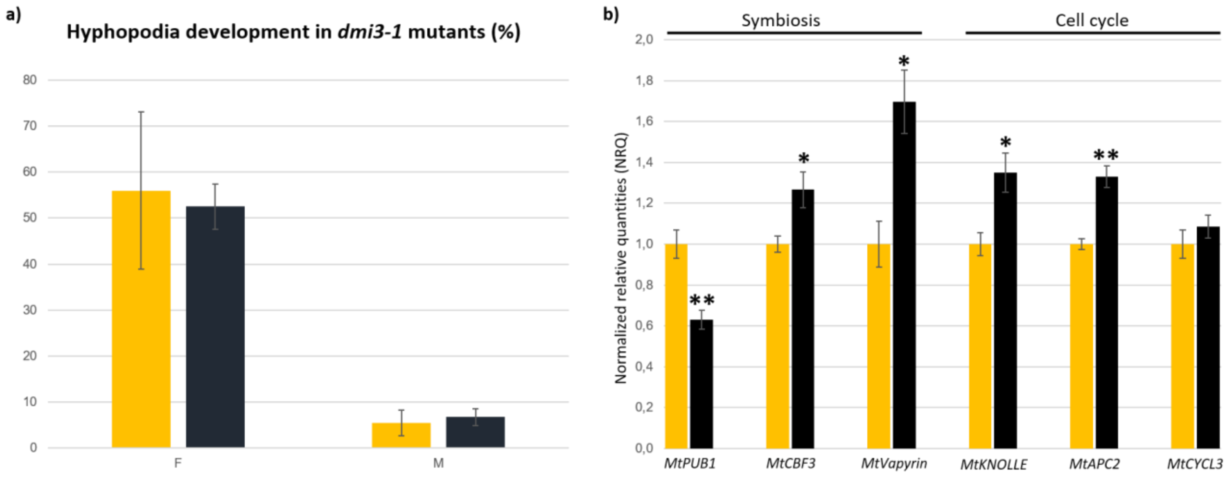


**Fig. S8 CO responses in *dmi3-1* mutants.** a) Hyphopodia assessment in mycorrhizal *dmi3-1* mutants, treated (black bars) or not (yellow bars) with CO. Both MYC and MYC+CO *dmi3-1* plants showed similar levels of hyphopodia. Histograms represent average values for each parameter for at least four biological replicates; error bars indicate standard deviation. b) Local CO treatment (1mg/ml) in *dmi3-1* ROCs stimulates the expression of different genes involved in fungal accommodation. Yellow bars = control, water treated roots; black bars = CO treated roots. Mean values ± SEs of four/six biological replicates for each treatment are shown. Asterisk indicate statistically significant difference (Student's t-test, *P < 0.05; **P < 0.001).

**Table S1 Summary statistics for the Illumina sequencing and mapping against *M. truncatula* genome assembly, 4.1 version.** Three biological replicates were analysed for each sample.

| Time point | Treatment | Sample ID | Rawreads (M) | % Aligned reads | Aligned reads (M) |
|---|---|---|---|---|---|
| 10 dpi | CTR | 1-C10-1 | 20.6 | 90.8% | 18.7 |
| | | 2-C10-2 | 26.6 | 90.6% | 24.1 |
| | | 3-C10-3 | 29.3 | 90.4% | 26.5 |
| | MYC | 4-M10-1 | 24.2 | 89.9% | 21.7 |
| | | 5-M10-4 | 28.9 | 90.6% | 26.2 |
| | | 6-M10-3 | 24.3 | 90.7% | 22.0 |
| | CTR + CO | 7-CCO10-1 | 24.6 | 91.7% | 22.6 |
| | | 8-CCO10-2 | 27.1 | 90.6% | 24.6 |
| | | 9-CCO10-3 | 28.8 | 91.1% | 26.3 |
| | MYC + CO | 10-MCO10-1 | 31.8 | 90.8% | 28.8 |
| | | 11-MCO10-3 | 21.5 | 90.9% | 19.6 |
| | | 12-MCO10-5 | 23.8 | 90.9% | 21.6 |
| 14 dpi | CTR | 13-C14-1 | 22.6 | 90.4% | 20.4 |
| | | 14-C14-2 | 19.2 | 90.9% | 17.4 |
| | | 15-C14-3 | 19.6 | 91.2% | 17.9 |
| | MYC | 16-CCO14-1 | 20.3 | 89.9% | 18.3 |
| | | 17-CCO14-2 | 20.9 | 90.2% | 18.8 |
| | | 18-CCO14-3 | 24.4 | 90.5% | 22.1 |
| | CTR + CO | 19-M14-1 | 25.8 | 91.3% | 23.5 |
| | | 20-M14-4 | 22.9 | 89.3% | 20.5 |
| | | 21-M14-3 | 29.7 | 89.0% | 26.4 |
| | MYC + CO | 22-MCO14-3 | 27.2 | 90.5% | 24.6 |
| | | 23-MCO14-4 | 19.4 | 90.5% | 17.6 |
| | | 24-MCO14-5 | 21.3 | 90.5% | 19.3 |
| 21 dpi | CTR | 25-C21-1 | 19.6 | 91.1% | 17.8 |
| | | 26-C21-2 | 22.9 | 91.0% | 20.9 |
| | | 27-C21-3 | 22.2 | 91.4% | 20.3 |
| | MYC | 28-CCO21-1 | 24.8 | 90.0% | 22.3 |
| | | 29-CCO21-2 | 21.6 | 90.5% | 19.5 |
| | | 30-CCO21-3 | 24.3 | 91.0% | 22.1 |
| | CTR + CO | 31-M21-4 | 24.8 | 88.8% | 22.0 |
| | | 32-M21-5 | 23.7 | 89.3% | 21.2 |
| | | 33-M21-3 | 30.0 | 87.2% | 26.2 |
| | MYC + CO | 34-MCO21-3 | 27.6 | 87.1% | 24.1 |
| | | 35-MCO21-4 | 27.8 | 89.0% | 24.7 |
| | | 36-MCO21-5 | 26.1 | 87.4% | 22.8 |
| 28 dpi | CTR | 37-C28-5 | 19.6 | 90.3% | 17.7 |
| | | 38-C28-2 | 26.4 | 91.1% | 24.1 |
| | | 39-C28-3 | 22.8 | 91.1% | 20.8 |
| | MYC | 40-CCO28-2 | 24.0 | 91.8% | 22.1 |
| | | 41-CCO28-1 | 22.4 | 91.5% | 20.5 |
| | | 42-CCO28-3 | 23.7 | 90.6% | 21.5 |
| | CTR + CO | 43-M28-3 | 22.3 | 90.3% | 20.1 |
| | | 44-M28-4 | 26.0 | 88.1% | 22.9 |
| | | 45-M28-5 | 19.4 | 89.3% | 17.3 |
| | MYC + CO | 46-MCO28-1 | 22.5 | 87.4% | 19.6 |
| | | 47-MCO28-3 | 19.8 | 88.8% | 17.6 |
| | | 48-MCO28-5 | 23.4 | 88.8% | 20.8 |

**Table S2** **List of over-represented MapMan functional categories.**

| BIN code | Description |
|---|---|
| 1 | Photosynthesis |
| 2 | Cellular respiration |
| 3 | Carbohydrate metabolism |
| 4 | Amino acid metabolism |
| 5 | Lipid metabolism |
| 6 | Nucleotide metabolism |
| 7 | Coenzyme metabolism |
| 8 | Polyamine metabolism |
| 9 | Secondary metabolism |
| 10 | Redox homeostasis |
| 11 | Phytohormone action |
| 12 | Chromatin organization |
| 13 | Cell cicle organization |
| 14 | DNA damage response |
| 15 | RNA biosynthesis |
| 16 | RNA processing |
| 17 | Protein biosynthesis |
| 18 | Protein modification |
| 19 | Protein homeostasis |
| 20 | Citoskeleton organozation |
| 21 | Cell wall organization |
| 22 | Vesicle trafficking |
| 23 | Protein translocation |
| 24 | Solute transport |
| 25 | Nutrient uptake |
| 26 | External stimuli response |
| 27 | Multi-process regulation |
| 35 | Not assigned-annotated |

**Table S3** List of primers used for RNA-seq validation experiments.

| Gene name | Primers sequence forward (5'-3') | Primers sequence reverse (5'-3') | Gene ID |
|---|---|---|---|
| *MtPUB1* | CATAGCCGAGGTAGGTGCTA | CCGAATTCGAGTACTTCGAC | Medtr5g083030 |
| *MtVAPYRIN* | GTTGGGTCAACCCCTCTTGA | CCATGTGTCCTTCTCTCGAA | Medtr6g027840 |
| *MtMYB1* | CTCTTTCCGAGGAATCTGAATC | GGTAGTGGATGATCCCAGTTC | Medtr7g068600 |
| *MtPT4* | TCGCGCGCCATGTTTGTTGT | GCGAAGAAGAATGTTAGCCC | Medtr1g028600 |
| *MtDMI2* | GGACGGGAACCTCTCAACAT | CCACAACTCTCCACAATGCCT | Medtr5g030920 |
| *MtWRKY* | GGGTGAGTCTTCAAAGCAACAAT | CCATCATCAAGAGGACCTTCCATT | Medtr3g090860 |
| *MtD14* | CCGCCGCTACACAACTCTTG | CACCTTGCTCAAATCCTCCGT | Medtr1g018320 |
| *MtCCD8* | GGGTCCATTCCTTTCCTGTTACTC | GCAGTCACCCTCCCATCTTCA | Medtr7g063800 |
| *MtCLE53* | TGTAGTGAGGTCAGAGGCAAGA | CCGCAGCCACTCACTATTTC | Medtr8g463700 |
| *MtTPP* | CATACTCCTCGCATTGCTGATACT | GCCAACCATTAGAACGAACATCAT | Medtr3g008500 |

**Table S4** **List of primers used for qRT-PCR experiments.**

| Gene name | Primers sequence forward (5'-3') | Primers sequence reverse (5'-3') | Gene ID |
|---|---|---|---|
| Primers used for gene expression analysis of strigolactones | | | |
| *MtDLK2* | GCAGTTCCAATAAGCATAGGGCAT | CTTCAGCACCTCAACCAACCTC | Medtr3g045440 |
| *MtMAX1a* | CGAAGAAATGAAACAAAGACACCC | TATGACACTGAACTTGACACCAT | Medtr3g104560 |
| *MtPDR1a* | CAGTGGAAGAGTTTGTTAGGAGTT | GCAAAGGCAGATCCAAAGATCAG | Medtr3g107870 |
| *MtCCD7* | CCAAGAATGTGATGTGAAGCC | TTTCTACTTCCAGCAGTCCAAG | Medtr7g045370 |
| *MtCCD8b* | GATCCTACAAGCTCTACCACTCAA | CACCTCCATCTCTAGGCACA | Medtr3g110195 |
| Primers used for gene expression analysis of early-signaling markers | | | |
| *MtPUB1* | CATAGCCGAGGTAGGTGCTA | CCGAATTCGAGTACTTCGAC | Medtr5g083030 |
| *MtCBF3* | CCAAAGAGGTTCCAAGAGATTAA | GCTATCTATCTACCATTCAACCCT | Medtr8g091720 |
| *MtGSTearly* | CTTGTTTACTTCCAATCTCGCAG | CACAAATACAAACGTACGCTACT | Medtr8g087425 |
| Primers used for gene expression analysis of fungal accomodation markers | | | |
| *MtKNOLLE* | CTCTCTGGACTTAAAGATGGTTCA | GCCTAAGTCCCTGAAACTCCAT | Medtr5g012010 |
| *MtAPC2* | TTTGCCCAGGACTTGATCCC | GAAGCTGGGTTCTGTCTGCT | Medtr2g084610 |
| *MtTPLATE* | GCATATTTGATACTAGGAGCG | CCCGTAAATTCTTCACAAGT | Medtr7g031450 |
| *MtCYCL2* | CACTTGCTCAGATAGAGTTCCC | GGTCATAAGGTAAAGGAGTTGAGG | Medtr2g118260 |
| *MtCYCL3* | CTGGAAGCTCCCTTTCCTTTA | CGATTTACCACTTCATTGTCTGAC | XM_003589009 |

**Dataset S1 (separate file).** RNA-seq based expression data for all genes expressed in CCO *vs* C during time-course experiment.

**Dataset S2 (separate file).** RNA-seq based expression data for all genes expressed in M *vs* C during time-course experiment.

**Dataset S3 (separate file).** RNA-seq based expression data for all genes expressed in MCO *vs* C during time-course experiment.

**Dataset S4 (separate file).** RNA-seq based expression data for all genes expressed in MCO *vs* M during time-course experiment.

**Dataset S5 (separate file).** Gene Ontology (GO) functional enrichments (Padj <0.1) of DEGs list in CTR+CO *vs* CTR.

**Dataset S6 (separate file).** Gene Ontology (GO) functional enrichments (Padj <0.1) of DEGs list in MYC *vs* CTR.

**Dataset S7 (separate file).** Gene Ontology (GO) functional enrichments (Padj <0.1) of DEGs list in MYC+CO *vs* CTR.

**Dataset S8 (separate file).** Gene Ontology (GO) functional enrichments (Padj <0.1) of DEGs list in MYC+CO *vs* MYC.
